# Life-cycle plasticity enables conditional asexual reproduction in a kelp

**DOI:** 10.64898/2026.06.18.732790

**Authors:** D Liesner, R Luthringer, L Kandjengo, JJ Bolton, MD Rothman, M Valero, FB Haas, SM Coelho

## Abstract

Many organisms alternate between distinct life stages and can reproduce either sexually or asexually, yet the mechanisms coordinating these developmental and reproductive transitions remain poorly understood. Here, we investigate the molecular basis of life stage identity and reproductive mode in the kelp *Laminaria pallida*, a multicellular organism with a complex life cycle. By combining gene expression analyses with developmental experiments and natural population-level approaches, we identify the genetic networks that govern developmental state transitions in both sexual and asexual contexts. We show that the capacity for asexual reproduction is broadly retained but rarely expressed in natural populations and is associated with reduced fitness. Moreover, we find that the presence of potential mating partners suppresses asexual development, revealing that reproductive mode is actively adjusted in response to reproductive opportunities. Together, these findings suggest that asexual reproduction functions as a conditional alternative to sex, promoting persistence when sexual reproduction is constrained while preserving the long-term benefits of sexual reproduction when mates are available. More broadly, our results reveal how organisms integrate developmental, environmental, and reproductive cues to balance alternative reproductive strategies across complex life cycles.

## Introduction

Sexual reproduction is widespread among eukaryotes and plays a central role in evolutionary processes by reshuffling parental genomes and generating new combinations of traits upon which natural selection can act [1]. In comparison with asexual reproduction, however, sexual reproduction entails considerable costs relating to, e.g., meiosis [2] or the production of males [3]. Sexuality has indeed independently been lost across many eukaryotic lineages [4,5]. While obligate asexuality is rare in animals and plants, other lineages such as fungi or many unicellular eukaryotes have never been observed to reproduce sexually [4]. The respective alternative life cycle strategies bypass meiosis and/or syngamy in processes such as parthenogenesis, apomixis, fragmentation or vegetative propagation [6,7]. Shifts between sexuality and asexuality may also occur within a single species and are often associated with life history trade-offs, developmental plasticity, or ecological adaptation [8,9], e.g., in geographical parthenogenesis at distributional range edges [10]. Facultative asexuals, which are able to switch between reproductive modes in a plastic response, offer a powerful system to explore the genetic and environmental controls governing reproductive mode transitions.

Brown algae (Phaeophyceae) provide a powerful system for studying reproductive mode switching because most species have complex life cycles and can reproduce both sexually and asexually [11–14]. In the brown alga *Scytosiphon*, female gametes develop parthenogenetically in natural populations, underscoring the ecological significance of asexual reproduction in this lineage [15]. This process is controlled by both the U sex chromosome and an autosomal locus [15]. A role for sex chromosomes in parthenogenesis has also been demonstrated in the model brown alga *Ectocarpus* [11]. In *Ectocarpus*, the sporophyte developmental program is activated in both zygotes and unfused gametes through the dimerization of two TALE homeodomain transcription factors [16,17], uncoupling life-cycle identity from ploidy. As a result, both male and female gametes can develop parthenogenetically into parthenosporophytes. Whether these parthenosporophytes consistently execute the diploid sporophyte developmental program, however, remains unknown.

Morphologically complex brown algae such as kelps also exhibit pronounced plasticity in their life cycles and reproductive modes. As ecologically important marine foundation species in diverse and highly productive coastal ecosystems [18], kelps display a marked contrast between their microscopic, filamentous gametophyte stage and their large, structurally complex sporophytes, which can reach several meters in length. Gametes are strongly dimorphic: small, motile sperm released by male gametophytes fertilize large, immotile eggs produced by females, initiating zygotic embryogenesis (**Figure 1A**). Notably, unfertilized female gametes (eggs) can undergo parthenogenetic development [19–21], although these parthenosporophytes are often morphologically abnormal and exhibit reduced fitness compared to sexually derived sporophytes [22–24]. Despite the potential ecological and evolutionary relevance, the molecular mechanisms governing reproductive mode transitions and plasticity in kelps, and particularly the regulation of gametophyte-to-sporophyte developmental switches remain largely unknown.

**Figure 1.**
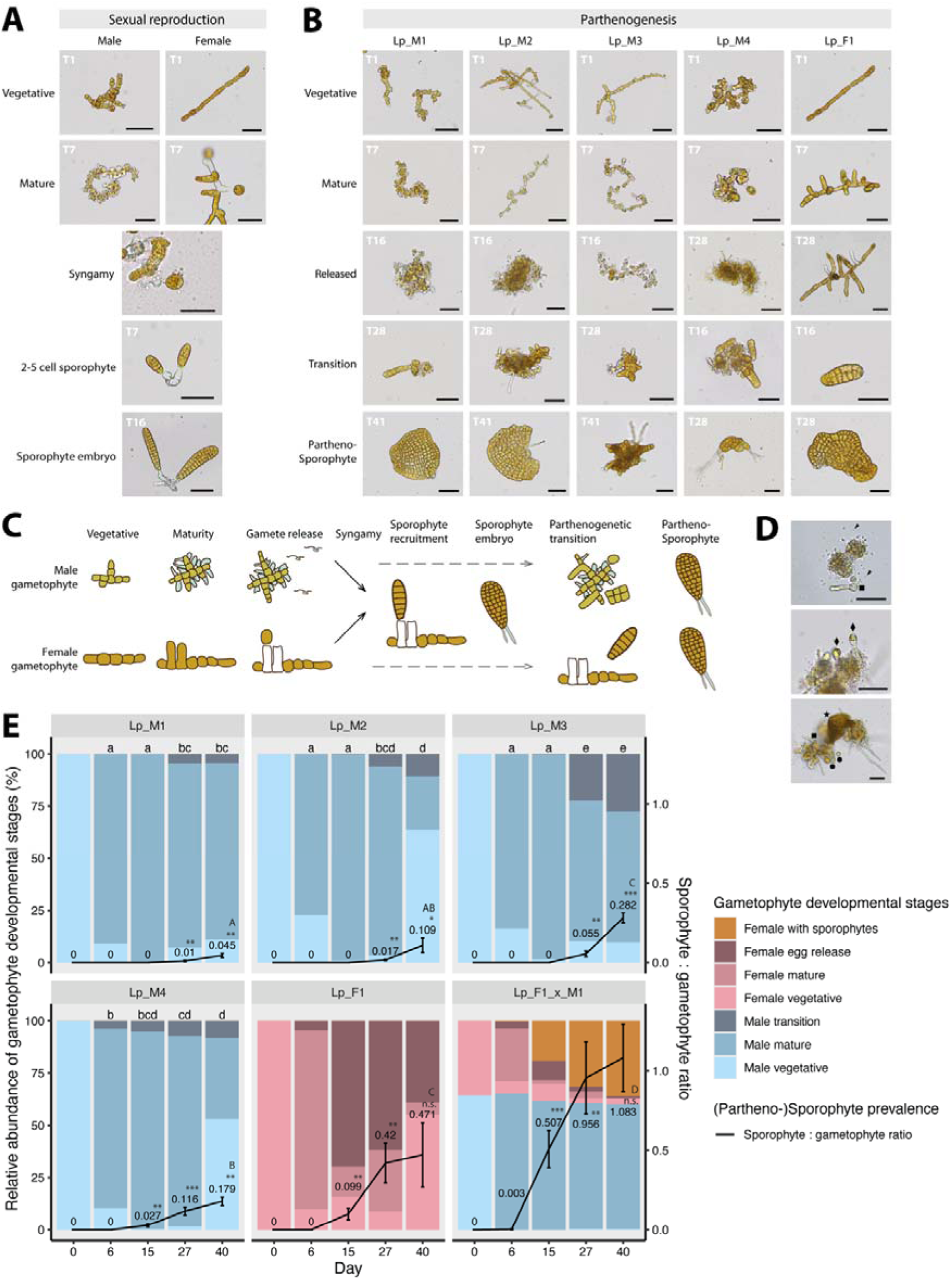
Developmental time course of *Laminaria pallida* gametogenesis and (partheno-)sporophyte production. **(A)** Three developmental stages were analysed during sexual reproduction of male and female gametophytes (vegetative, mature, female egg) and two stages of sporophyte embryo development (2-5 cell sporophyte, sporophyte embryo). **(B)** Five developmental stages during parthenogenetic development were analysed (vegetative, mature, after gamete release, morphological transition / early partheno-sporophyte tissue development, partheno-sporophyte) and collected at five time points (T1, T7, T16, T28, T41). Parthenogenesis occurred in all four male gametophytes (Lp_M1 – Lp_M4) and in the female Lp_F1. Note that gametophytes of Lp_M1 which did not transition were used as a control for all time points. Additionally, sporophytes developed from sexual reproduction of Lp_F1 and Lp_M1 were used as comparisons. **(C)** Schematic depiction of sexual and parthenogenetic reproduction in *L. pallida*, partially modified from [28]. **(D)** In addition to sperm (arrowheads) male gametophytes produce enlarged gametangia (diamonds) which release egg-like gametes (circles) which “germinate” (squares) and grow into partheno-sporophytes (star). **(E)** Relative abundance of partheno-sporophytes in cultures of four male strains as compared to a female line and to the presence of sexually derived sporophytes. Average relative abundance of developmental stages over time is shown in stacked bar plots (n = 5 replicate dishes per strain; primary y-axis). Significant differences in male ‘transition’ phenotypes across time and lineages are indicated by lowercase letters (linear model; Tukey post-hoc test; p < 0.05). Prevalence of (partheno-)sporophytes normalized to total number of gametophytes is shown as black line and text overlay (n=5, mean ± SD). Significant increases in (partheno-)sporophyte numbers per time step within gametophyte lineages are indicated by asterisks (first asterisk, T-test; following asterisks, linear model; Tukey post-hoc tests ***, p < 0.001; **, p < 0.01; * p < 0.05; n.s., not significant). Significant differences in sporophyte prevalence across gametophyte lineages on day 40 are indicated by capital letters (linear model; Tukey post-hoc tests; p < 0.05). All scale bars = 50 µm. Note that for simplicity, all linear model results are collated in **Supplementary Table S2**.

Here, we use the kelp *Laminaria pallida* as a model system to investigate the regulation of developmental plasticity. By combining transcriptomic analyses of sexual and asexual developmental pathways with population-level surveys across the species’ natural range, we characterize the molecular and ecological factors that govern developmental fate decisions. We show that gametes of *L. pallida* initiate sporophyte development without fertilisation through parthenogenesis. We characterise core gene expression programs shared along development between asexual and sexual reproduction. Although both male and female gametes can develop parthenogenetically under laboratory conditions, extensive screening of natural populations revealed no evidence that asexual reproduction contributes substantially to population maintenance in the wild. We demonstrate that the presence of the opposite sex inhibits parthenogenetic development, suggesting that a diffusible signal promotes sexual reproduction and limits alternative asexual developmental trajectories. Together with the reduced fitness of parthenogenetically derived sporophytes, this regulatory mechanism likely explains why asexual development remains rare in nature despite its developmental feasibility. These findings reveal an unexpected degree of developmental flexibility in a major marine lineage and provide new insights into how reproductive mode, developmental regulation, and ecological context interact to shape life-cycle evolution. More broadly, they advance our understanding of the mechanisms that maintain sexual reproduction despite the widespread latent capacity for asexual development and have implications for the conservation, restoration, and cultivation of kelp forests in a changing ocean.

## Results

### Parthenogenetic development is not restricted to female gametes

Parthenogenetic development of female gametes has been described in several kelp species, but whether male gametes can similarly initiate sporophyte development remains unclear [22,25,26]. To compare sexual and asexual developmental trajectories, we compared sporophyte formation in mixed and unmixed male and female gametophyte cultures (**Table S1; Figure 1A,B**).

In unmixed cultures, we identified five distinct developmental stages: (1) vegetative gametophytes; (2) mature gametophytes bearing gametangia; (3) gametophytes after gamete release; (4) a transitional stage marked by the initiation of parthenosporophyte development; and (5) early parthenosporophyte formation, characterised by the emergence of two-dimensional blades (**Figure 1B,C**). Importantly, direct observations confirmed that male parthenosporophytes originated from individual sperm cells (**Figure 1D**; [27]), revealing no evidence for apogamy (i.e., development of sporophytes directly from vegetative gametophytes).

We next quantified the prevalence and timing of parthenogenetic development (**Figure 1E**). In all male lines, a subset of gametophytes underwent a developmental transition towards parthenosporophyte formation, characterised by the release of enlarged gametes from gametangia with an oogonium-like morphology (**Figure 1D**). This transition occurred during or shortly after gametangial maturation, between days 6 and 27 (**Figure 1E**). By day 40, the proportion of male gametophytes undergoing a parthenogenetic transition ranged from 4.22 ± 1.13% (Lp_M1) to 27.72 ± 1.71% (Lp_M3), with the latter not significantly different from the female rate (Tukey test, p = 0.3492). Importantly, despite the capacity for parthenogenesis in both sexes, sexual reproduction was substantially more efficient. Co-cultivation of male and female gametophytes produced more than twice as many sporophytes as parthenogenesis alone (Tukey test, p < 0.01), with an average of one sporophyte embryo generated per gametophyte.

Together, these results demonstrate that both female and male gametes can initiate sporophyte development in the absence of fertilisation. Although the frequency of male parthenogenesis varied among strains, some male lines produced partheno-sporophytes at rates comparable to females. Sexual reproduction nevertheless remained substantially more effective in generating sporophytes.

### Partheno-sporophytes exhibit abnormal development, endoreduplication, and reduced fitness

A subset of male and female partheno-sporophytes, as well as sexually derived sporophytes, were isolated and cultured for further development (see Methods). After 31 weeks, clear morphological differences were observed between partheno-sporophytes and diploid sporophytes (**Figure 2A**). Sporophytes exhibited well-defined apical–basal thallus organisation, including differentiated holdfast, stipe, and blade regions. In contrast, parthenosporophytes displayed developmental abnormalities, such as malformed blades and occasionally incomplete differentiation of key tissues, particularly the holdfast and stipe. Extensive lateral growth often led to folds in the blade of partheno-sporophytes. Despite these defects, parthenosporophytes could be grown for several months but remained significantly smaller compared to sexually derived sporophytes (Kruskal Wallis Χ^2^=27.78, p=1.36*10^−7^; **Figure 2B**). PCR sex markers confirmed the presence of only one sex chromosome in asexually produced parthenosporophyte tissue (**Figure 2C**) and ploidy analysis by flow cytometry indicated that endoreduplication had occurred in all tested partheno-sporophytes (**Figure S1**). Despite three independent induction attempts, parthenosporophytes remained infertile under laboratory conditions.

**Figure 2.**
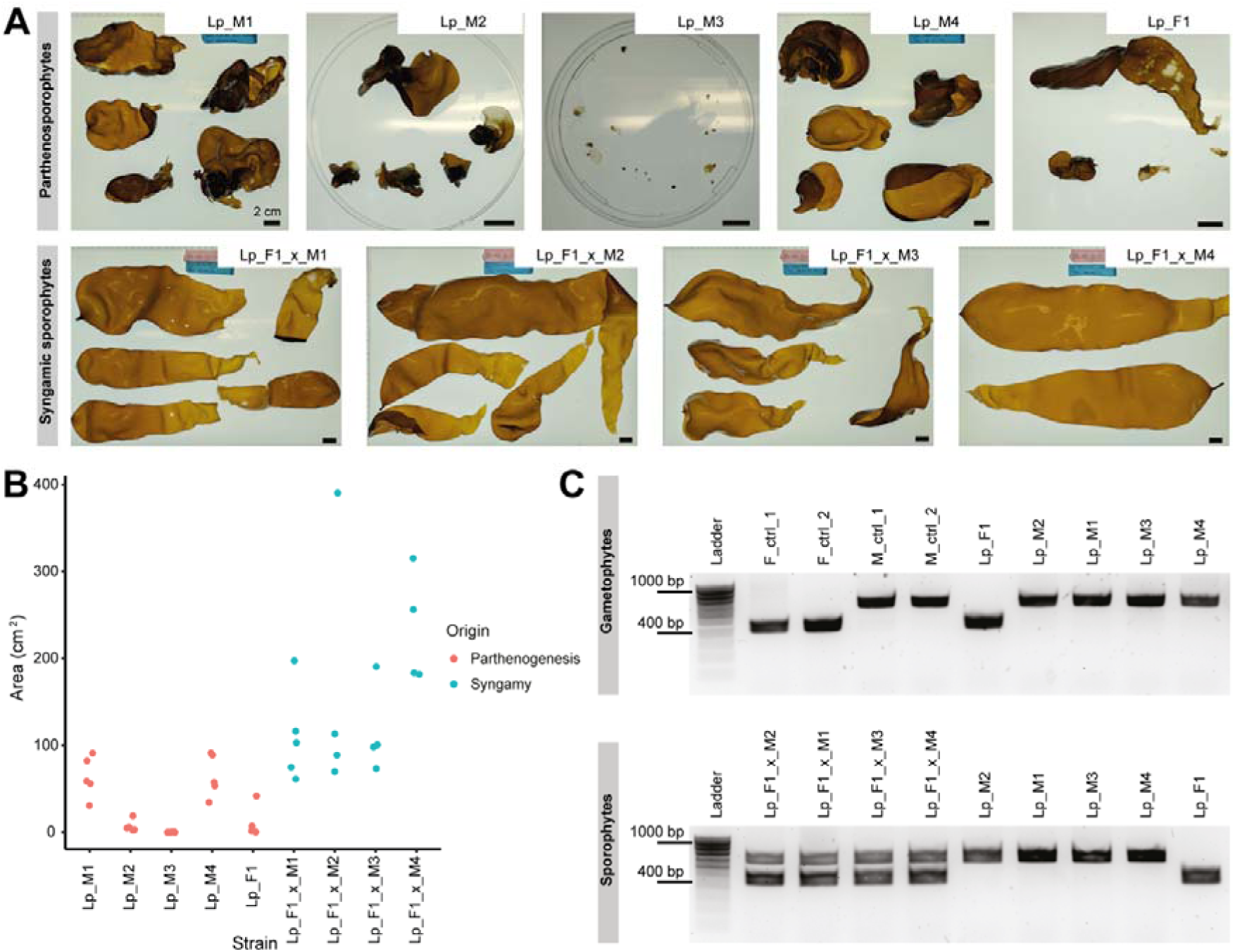
*Laminaria pallida* sporophyte characterisation. **(A)** Morphology of parthenogenetically and sexually derived sporophytes. Scale bar = 2 cm. **(B)** Sporophyte surface area of partheno-sporophytes (red) and sexually derived sporophytes (blue). **(C)** Gel electrophoresis of sex-specific PCR markers indicating presence of a U-chromosome (450 bp) and a V-chromosome (700 bp) in gametophytes, partheno-sporophytes and sexually derived sporophytes.

Together, these results indicate that partheno-sporophytes may arise from both male and female gametes in kelp, that their development involves endoreduplication, and that they exhibit substantially reduced fitness compared to diploid sporophytes formed via sexual reproduction.

### Convergence of transcriptional signatures during parthenogenesis

To identify the molecular changes underlying the onset of parthenogenetic development and compare them with those associated with zygotic embryogenesis, we performed RNA sequencing (RNAseq) across defined developmental stages (**Figure 1A,B**). Principal component analysis (PCA) revealed distinct transcriptional signatures associated with sex and developmental stage (**Figure 3A**). Notably, although male gametophytes underwent pronounced transcriptomic changes upon reaching fertility, transcriptional trajectories from male and female parthenogenesis and sexual reproduction converged during the transition to sporophyte development.

**Figure 3.**
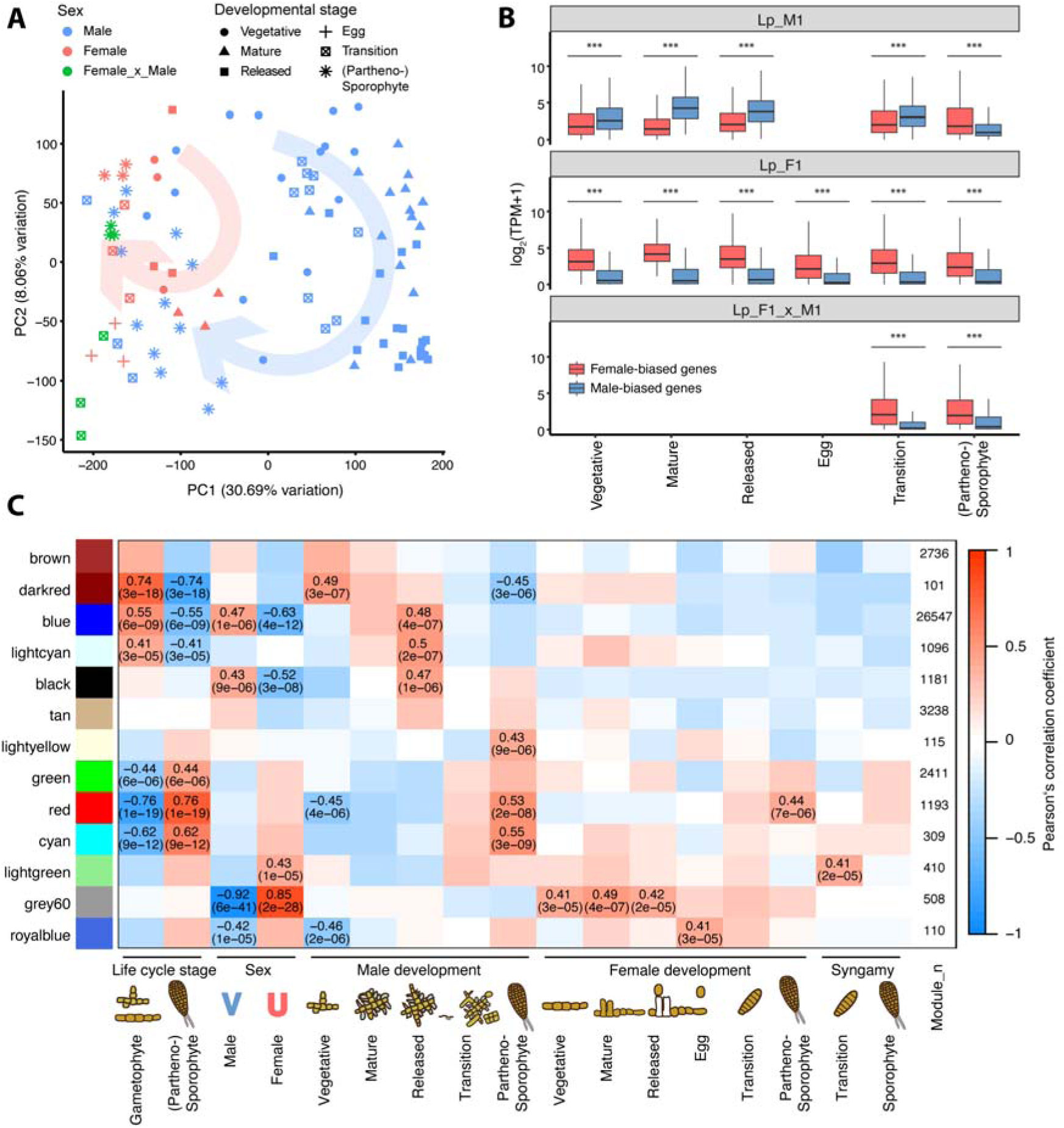
Gene expression patterns correlated to phenotype expression in *Laminaria pallida*. **(A)** Principal component analysis of RNA sequencing libraries with genetic sex indicated by colour and symbols distinguishing developmental stages (n=98). Arrows indicate transcriptomic convergence along development towards sporophyte identity. (B) Sex-biased gene expression for male Lp_M1, female Lp_F1 and cross Lp_x_M1 across developmental stages; outliers are excluded for clarity; significant differences in sex-biased gene expression are indicated for each developmental stage (Wilcoxon rank sum test; ***, p < 0.001). **(C)** Relation of module eigengenes to life cycle stages, genetic sex, and developmental stages of gametophyte and (partheno-) sporophyte development. For each module and trait, the heatmap indicates Pearson’s correlation coefficient (R), and for |R| > 0.4, both R and p-value are displayed. Schematic drawings are partially modified from [28].

Sex-biased genes were identified for each strain as differentially expressed genes between male and female gametophytes during maturity (DESeq2; |log2FC| > 1; p < 0.001). While male-biased genes were dominantly expressed throughout development in all male strains, a significant reduction of male-biased gene expression occurred during partheno-sporophyte development (Wilcoxon tests, p < 0.0001; **Figure 3B; Figure S3**), leading to a transcriptomic de-masculinisation that resulted in expression profiles resembling those of female partheno-sporophytes and zygotic sporophyte embryos.

Given the convergence of male partheno-sporophyte and zygotic embryonic transcriptomes, we next examined the expression of conserved regulators of the gametophyte-to-sporophyte transition [17]. Two TALE homeodomain (HD) transcription factors that govern this developmental switch in the model brown alga *Ectocarpus*, SAMSARA and OUROBOROS [16,17], were identified in the *L. pallida* genome based on sequence homology (**Table S3; Figure S4**). Neither gene exhibited significant stage-specific expression (Wilcoxon rank sum tests, p > 0.05; no consistent differential expression, DESeq2). Instead, both were expressed throughout gametophyte and partheno-sporophyte development.

We identified a core set of transcripts that was differentially expressed between partheno-sporophytes and vegetative gametophytes of all lineages. Overall, 222 DEGs were consistently up-regulated and 84 transcripts were consistently down-regulated (**Figure S5; Table S3**). Homology search against the *Ectocarpus* genome revealed that transcripts related to known life cycle regulators were commonly up-regulated, namely three transcripts for *imm upregulated 13*, along with genes related to photosynthesis (light harvesting complex), cell wall metabolism (mannuronan C-5-epimerase), and brown algal viruses (EsV-1-7 domain).

A weighted gene co-expression network analysis (WGCNA) highlighted several co-expression modules related to differentiation between sexes and life cycle stages (**Figure 3C**). Notably, the largest module blue comprised 66.4% of all transcripts and was significantly associated with male gametophyte sexual development. This association, however, ceased during the transition stage and became significantly negative during partheno-sporophyte development, indicating de-masculinisation of the transcriptome. The strongly enriched biological process gene ontology (GO) term “microtubule-based movement” indicates involvement in male gamete production (**Figure S6**). Module grey60 was continuously associated with female development with a peak during maturity (**Figure 3C**), and was strongly enriched in the molecular function “DNA binding” (**Figure S7**). Modules red and cyan were both significantly correlated to partheno-sporophyte development (**Figure 3C**). In addition to metabolic processes related to photosynthesis and energy metabolism (“ATP metabolic process”, “lipid metabolic process”, “photosynthesis”, “light harvesting”), the enrichment of “peroxidase activity” and “oxidoreductase activity” indicates ROS signalling involved in sporophytic development (**Figures S8, S9**). In module cyan, which is only significantly related to male but not female parthenogenesis (**Figure 3C**), the enrichment of “box H/ACA snoRNP assembly” and “protein folding” may be related to post-transcriptional and post-translational modifications specific to male parthenogenesis.

Together, these results indicate that male parthenogenesis is accompanied by transcriptome de-masculinisation, leading to convergence with female parthenogenesis and zygotic embryo development. Nevertheless, distinct expression signatures in life cycle–related genes and large co-expression modules suggest that these developmental trajectories remain molecularly distinct.

### No evidence for parthenogenetic reproduction in natural populations

To assess whether parthenogenesis has a relevant contribution to population structure and diversity in natural populations of *L. pallida*, we sampled sporophyte tissue from the populations of Paternoster, South Africa (ZAF) and Swakopmund, Namibia (NAM; **Figure S2A; Table S4**). At each site, we employed a hierarchical sampling design spanning spatial scales from a few metres to several hundred metres. Specifically, we sampled at least 20 individuals from each of eight areas separated by <5 m within each population (**Figure 4A; Figure S2B**). Using ten microsatellite markers (see Methods), we detected significant genetic differentiation between the ZAF and NAM populations (FST = 0.293, p = 0.050). Structure analysis identified K = 2 as the most likely number of genetic clusters following Evanno et al. [29] (**Figure S2D**), corresponding to the two sampled populations. No additional genetic substructure was detected within either population at K ≥ 3. Estimates of the inbreeding coefficient (F_IS_) did not differ significantly from expectations under random mating in either population (**Figure 4B**; **Table 1**), and F_IS_ did not differ between ZAF and NAM (F_1,18_ = 1.613, p = 0.22). Moreover, F_IS_ remained consistent across spatial scales within each population (**Figure S2C).** Together, these results indicate random mating and an absence of detectable fine-scale spatial genetic structure within the two populations.

**Figure 4.**
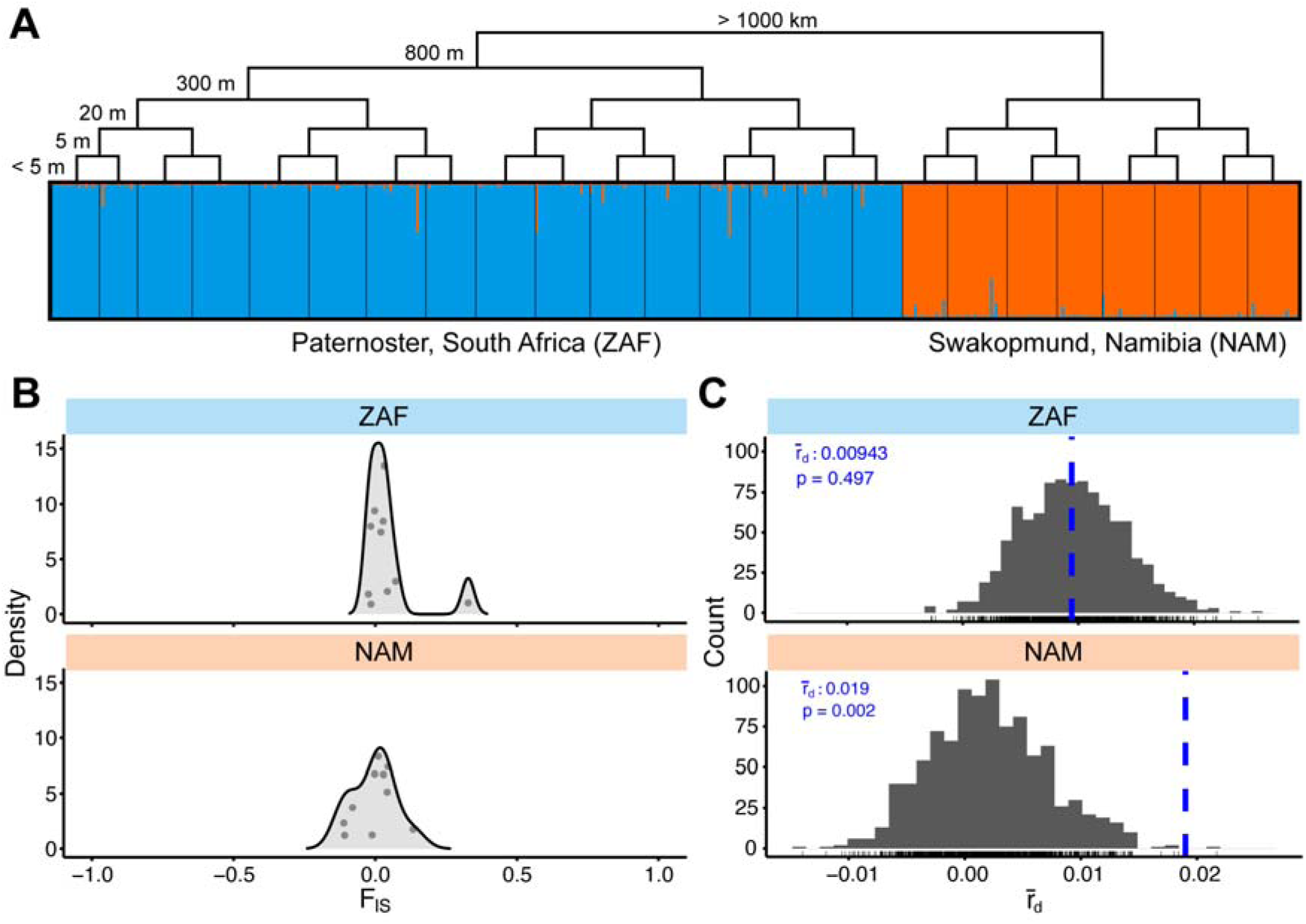
Population genetic characteristics of *Laminaria pallida* sampled at Paternoster, South Africa (ZAF; n=357) and Swakopmund, Namibia (NAM; n=166). **(A)** Structure bar plot obtained for K = 2 genetic clusters. Tree reflects the hierarchical sampling levels in decreasing order of magnitude. Individuals (vertical bars) were assigned probabilities of belonging to clusters (colors) based on differences in genetic variance. **(B)** Density plot of inbreeding coefficient F_IS_ for all microsatellite loci (n=10) in ZAF and NAM. **(C)** Test for linkage disequilibrium by random permutations (n=999) of the standardised index of association (rlJd) shown as histograms and observed values shown by dashed blue lines for NAM and ZAF.

**Table 1.**
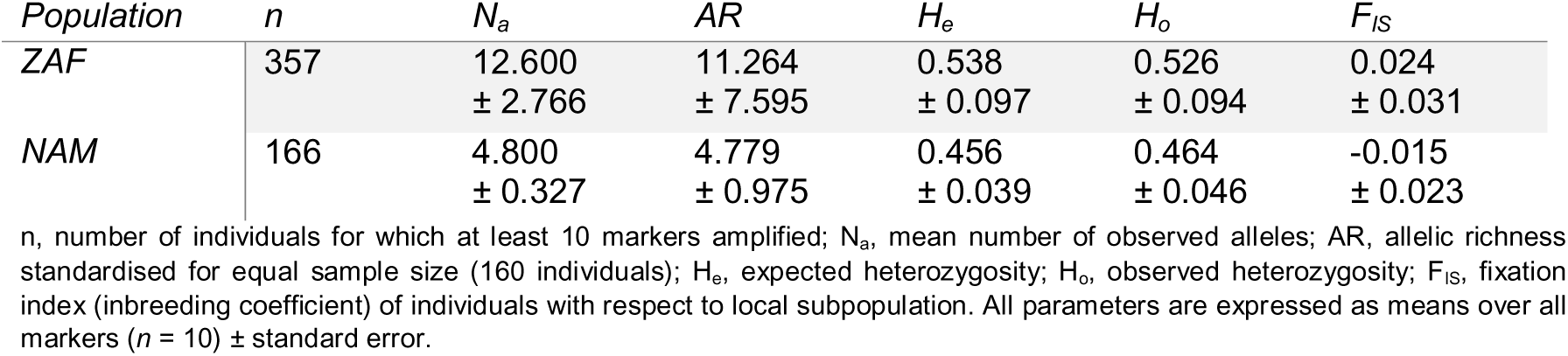
Genetic characteristics of the distinct *Laminaria pallida* populations of Paternoster, South Africa (ZAF) and Swakopmund, Namibia (NAM).

The number of alleles N_a_ was significantly higher in ZAF than in NAM (ANOVA; F_1,18_ = 7.84, *p* = 0.012; **Table 1**), as was the allelic richness AR (ANOVA; F_1,18_ = 6.45, *p* = 0.021). However, neither expected heterozygosity H_e_ (ANOVA; F_1,18_ = 0.623, *p* = 0.44) nor observed heterozygosity H_o_ (F_1,18_ = 1.90, *p* = 0.185) differed significantly between populations. Additionally, we did not identify unambiguous evidence of parthenogenesis among the genotyped individuals, as no identical multilocus genotypes and no purely homozygous genotypes were identified. However, analysis of linkage disequilibrium based on the standardised multilocus index of association rld revealed a significant deviation from random mating only in NAM but not in ZAF (**Figure 4C**). Together, these results indicate that asexual reproduction and clonality is not prevalent in nature.

To assess whether wild populations retain the capacity for asexual development despite its apparent absence in the field, we established 31 female and 28 male gametophyte cultures from a subset of collected sporophytes. Although bacterial contamination affected few samples and six isolates did not become reproductive, 92% of mature male and female gametophytes produced partheno-sporophytes (**Table S5**). These results indicate that the potential for clonal reproduction is broadly maintained in both sexes within natural populations, even though it may be ecologically or competitively disadvantaged compared to sexual reproduction.

### Male parthenogenesis is suppressed in the presence of females

The analyses described above revealed no evidence of clonal reproduction in natural populations. This raises the question of how the capacity for parthenogenesis is maintained despite its apparent absence in natural populations. One possibility is that parthenogenesis provides reproductive assurance when mates are unavailable. To test this hypothesis, we examined whether the presence of a mating partner in the same culture medium influences the frequency of parthenogenetic development (see Methods). If so, parthenogenesis could function as a conditional reproductive strategy that is activated under mate limitation.

To test this hypothesis, we used the two male lines with the highest rates of parthenogenesis (Lp_M3 and Lp_M4; **Figure 1E**) together with the female line Lp_F1. In both male lines, the frequencies of transition phenotypes (**Figure 5A**) and partheno-sporophyte development (**Figure 5B**) were significantly higher in the absence of a female partner, indicating that mate presence suppresses parthenogenesis. Females showed a similar response, producing more egg-stage gametophytes and partheno-sporophytes when males were absent. Moreover, females released significantly more eggs in the absence than in the presence of males (2.61 ± 0.39 versus 1.75 ± 0.12 eggs per individual on day 6; t-test, t = 4.677, p = 0.0062). Together, these results reveal a bidirectional mate-recognition mechanism that promotes sexual reproduction while actively suppressing parthenogenetic development in both sexes.

**Figure 5.**
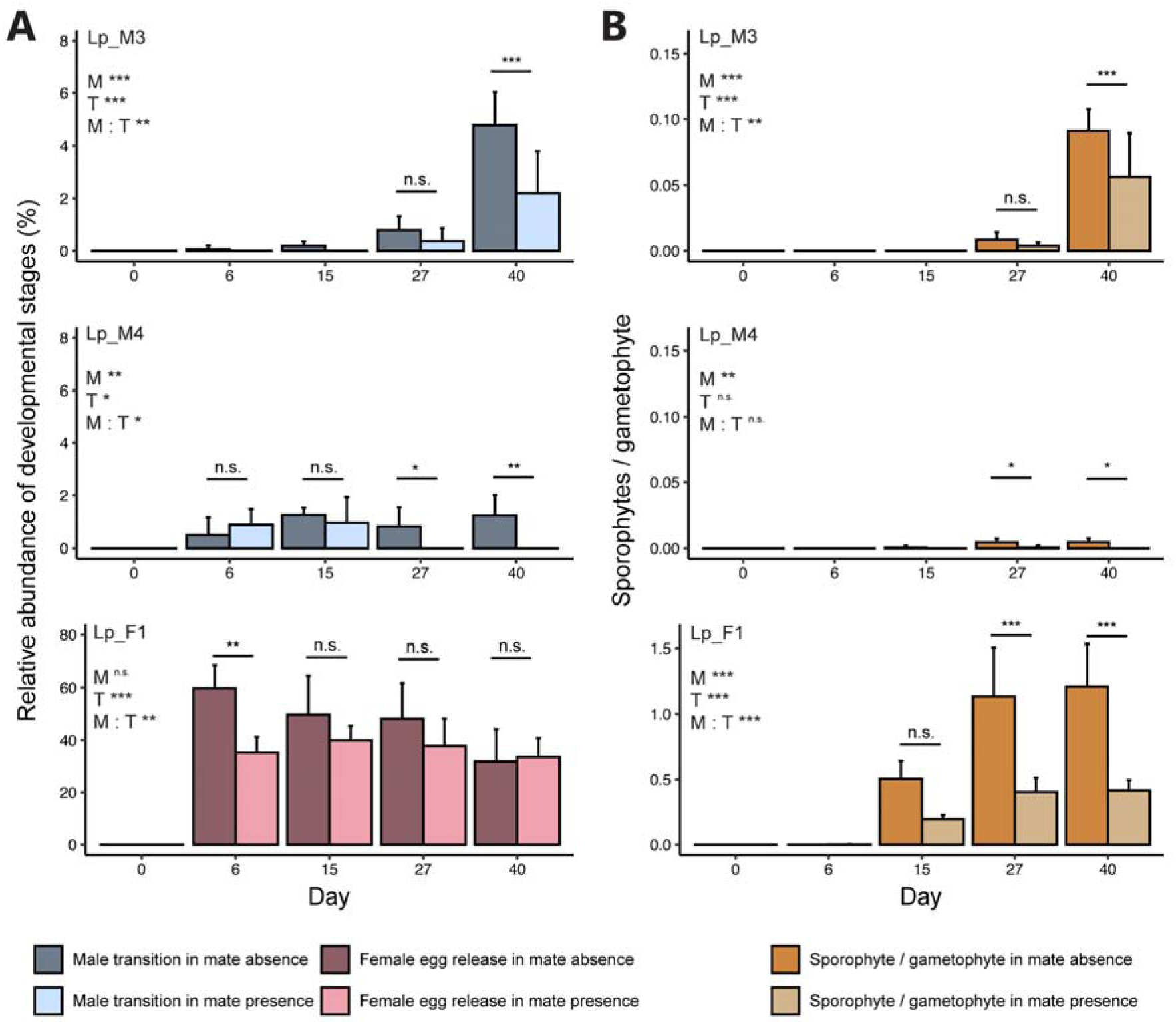
Effect of mate presence over time on the prevalence of parthenogenesis in *Laminaria pallida* gametophyte males (Lp_M3; Lp_M4) and females (Lp_F1) with respect to **(A)** gametophyte development and **(B)** sporophyte production. Statistical significance of the fixed factors mate presence (M), time (T), and their interaction (M : T), and results of pairwise comparisons (Tukey test) are summarised in the figure; ***, p < 0.001; **, p < 0.01; * p < 0.05; n.s., not significant. Note that for simplicity, all linear model results are collated in **Supplementary Table S6**.

## Discussion

### Mate-dependent regulation of parthenogenesis in kelps

Our results reveal an unexpectedly broad capacity for asexual reproduction in the kelp *Laminaria pallida*. All gametophyte lines tested were capable of parthenogenetic development under laboratory conditions. This trait is unlikely to be unique to *L. pallida*, as female parthenogenesis is widespread across kelps and male parthenogenesis has been reported from several genera worldwide [22,25,26,30], with growing evidence that asexual development may be more common than previously appreciated [27,31,32].

Despite this latent capacity, sexual reproduction clearly outperformed parthenogenesis in our experiments. Male parthenogenesis was slower and produced fewer offspring than female parthenogenesis, while sexually derived offspring were both more abundant and grew more rapidly than progeny produced through either male or female parthenogenesis. These findings indicate that sexual reproduction retains a substantial fitness advantage whenever mates are available.

Parthenogenesis is common among morphologically simpler brown algae [11,15], whereas more complex lineages such as fucoids and kelps are generally thought to rely on sporophyte-mediated forms of clonal propagation. In *Fucus vesiculosus*, for example, extensive single-sex clonal populations have become established in the Baltic Sea through fragmentation and reattachment [33,34]. Likewise, several kelp species propagate vegetatively through clonal blade production from creeping stolons [35–37]. Nevertheless, female partheno-sporophytes have occasionally been reported to develop normally [38] and even reach reproductive maturity in natural populations [30], suggesting that gamete-derived asexual reproduction remains biologically relevant in kelps.

Yet, despite the universal capacity for parthenogenesis observed in our experiments, we found no evidence for clonal reproduction in natural populations of *L. pallida*. This apparent paradox is resolved by our mate-exclusion experiments, which demonstrate that parthenogenesis is actively regulated by mate availability. In both sexes, parthenogenetic development increased when mating partners were absent and declined when they were present. These findings point to a mate-recognition mechanism that promotes sexual reproduction while suppressing asexual development. Rather than representing a competing reproductive strategy, parthenogenesis may therefore function as a conditional backup pathway that provides reproductive assurance when mates become limiting, while remaining largely masked under conditions that favour sexual reproduction.

### Shared developmental programs underpin parthenogenesis and embryogenesis

In *Ectocarpus*, the two TALE homeodomain transcription factors that trigger sporophyte development are present in both sexes, rendering life-cycle identity independent of syngamy [11,17,28]. Our results suggest that kelp gametophytes similarly possess the molecular machinery required to initiate the sporophyte program autonomously. However, unlike in *Ectocarpus*, both *SAMSARA* and *OUROBOROS* are continuously expressed throughout male and female development in *Laminaria pallida*, suggesting that regulation of these key life-cycle regulators differs between the two species[16,17].

We identified core sets of genes and co-expression modules associated with sporophyte development during both parthenogenesis and zygotic embryogenesis. Among the genes shared across developmental trajectories were homologs of genes controlled by *IMMEDIATE UPRIGHT* (*imm upregulated 13*), which controls the initial asymmetric cell division during sporophyte development in *Ectocarpus* [39,40]. Reactive oxygen species (ROS), which play key roles in embryogenesis in both plants and brown algae [41,42], were also strongly represented among parthenogenesis-associated modules. These findings suggest that parthenogenetic and zygotic development recruit a common developmental program, despite their distinct origins. Because transitions between sexual and asexual reproduction can involve diverse genetic mechanisms [43], future studies will be required to identify the loci that regulate parthenogenesis in kelps.

Against this shared developmental background, male parthenogenesis is accompanied by a striking transcriptional reconfiguration. We previously showed that reduced expression of male-biased genes results in a feminized gametophyte morphology, whereas male development depends on a complex network of sex-biased gene expression [28]. Consistent with this, male development in both *Macrocystis pyrifera* and *L. pallida* is associated with expression of a large fraction of the transcriptome [28]. Here, we show that parthenogenetic development is preceded by extensive de-masculinisation of the transcriptome following gamete release. Concomitantly, male gametes enlarge, lose the capacity for fusion, and acquire the ability to initiate parthenogenesis (see also [27]). Together, these observations suggest that male parthenogenesis involves a partial reprogramming away from a canonical gametic state, generating cells that may function primarily as clonal dispersal propagules rather than as gametes *sensu stricto*.

### Parthenogenesis as a conditional reproductive strategy

Despite the widespread occurrence of parthenogenesis in our laboratory experiments, we detected no unambiguous population-genetic signatures of asexual reproduction among 523 sporophytes sampled from Namibian and South African populations. This discrepancy suggests that parthenogenetically derived offspring contribute little to recruitment in natural populations. One explanation is that asexual offspring experience reduced fitness relative to sexually derived sporophytes, consistent with the delayed development and growth defects observed here. Population genetic modelling in giant kelp similarly predicts high early mortality among highly homozygous offspring produced through self-fertilisation [44].

Our experiments further indicate that parthenogenesis is actively regulated by mate availability. In both sexes, asexual development increased when mating partners were absent, suggesting that a mate-recognition mechanism promotes sexual reproduction while suppressing alternative developmental trajectories. Similar patterns have been proposed in giant kelp, where female egg production increases in the absence of males [46], and are consistent with the hypothesis that parthenogenesis functions as a form of reproductive assurance under mate limitation [45,46]. Such conditions may occur in sparsely populated habitats or during episodes of environmental stress, potentially facilitating persistence at distributional margins [10].

Evidence for such a role in natural populations, however, remains limited. We detected a weak but significant deviation from random mating in the Namibian population that could reflect cryptic clonality. Whether this pattern is related to local environmental conditions remains unclear. Sea surface temperatures at this site regularly exceed 20°C (**Figure S2E**), a threshold that may impair growth and reproduction in *L. pallida* [47], potentially increasing the relative importance of asexual developmental pathways.

Together, these observations suggest that the evolutionary significance of parthenogenesis in kelps is likely to depend on ecological context. There is currently no evidence that parthenogenesis becomes more frequent at distributional range edges, despite reports of reproductive abnormalities in populations of *Laminaria digitata* near their warm range limit [48]. Moreover, parthenogenetically derived sporophytes appear less competitive than their sexually produced counterparts, owing to delayed development, irregular growth, and, in some species, reduced fertility [49]. Yet the evolutionary maintenance of this trait may not require substantial fitness benefits. Male gametophytes produce functional sperm [27], and in our experiments most partheno-sporophytes developed only after the main period of gamete release. Thus, investment in asexual development may occur largely after opportunities for sexual reproduction have been exhausted. If so, parthenogenesis may persist because it incurs little additional cost while providing a low-probability route to reproduction when sexual reproduction fails.

In conclusion, our results reveal a widespread but tightly regulated capacity for parthenogenesis in kelps. Parthenogenetic and sexual development converge on a common sporophyte program, yet mate availability biases reproduction towards sex, potentially restricting the contribution of asexual offspring in natural populations. These findings suggest that parthenogenesis functions not as an alternative to sexual reproduction, but as a latent developmental pathway that may provide reproductive assurance when opportunities for mating are limited.

## Materials and Methods

### Study material

Fertile sporophytes of *Laminaria pallida* had been collected on the shores of Kommetjie, South Africa, in 1999 (Lp_M1–Lp_M3), Swakopmund, Namibia, in 1993 (Lp_M4), and Paternoster, South Africa in 2022 (Lp_F1). Meiospores were released and clonal gametophyte isolates were prepared as described by [50]. All gametophytes were grown vegetatively in 50% Provasoli-enriched natural seawater (PES; [51]; iodine enrichment following [52]) under red light in a 14:10 h light:dark (L:D) cycle (MaxLED 500 RGBW, Paulmann, Springe, Germany) at 14°C in a thermostatic cabinet (TC 445 L, Lovibond, Dortmund, Germany).

### Culture conditions

To induce gametophyte fertility, gametophyte tufts were carefully ground using mortar and pestle, sieved to obtain a size fraction of 50-100 µm, and sowed into 55 mm petri dishes into 50% PES to a final volume of 12 mL. After a recovery period of two days at reduced irradiance (5-10 µmol phot. m^−2^ s^−1^), dishes were cultivated at 12°C in a 16:8 h L:D cycle at 20-30 µmol phot. m^−2^ s^−1^ white light. Once per week, 5 mL of culture medium were replenished with 50% PES.

Sporophytes obtained from syngamy and asexually were isolated with tweezers at a size of ∼1 mm. First, sporophytes were grown in 1L flasks with gentle aeration in a temperature-controlled chamber at MPI Tübingen at 14°C in a 12:12 h L:D cycle at 30–40 µmol photons m^−2^ s^−1^ in 100% PES. After four months, they were transferred to the Station Biologique de Roscoff, France, and grown at 13 ± 1°C in a 16:8 h L:D cycle at 30–40 µmol photons m^−2^ s^−1^ (T8 Lumilux 36W–865 Cool Daylight, Osram) in 100% PES. With increasing sporophyte size, culture volume was sequentially increased to 10L transparent round bottles (Nalgene). Medium was changed weekly, photos were taken on an LED table and sporophyte area was analysed using ImageJ Fiji [53].

### DNA isolation and sex marker PCR

To determine the genetic sex of gametophytes, partheno-sporophytes and crossed sporophytes, genomic DNA was extracted from 50-100 mg tissue using the OmniPrep Genomic DNA isolation kit (G-Biosciences, St. Louis, USA) following the manufacturer’s instructions. For sex genotyping, we used the male marker ldig-hmg-g1-3-4 (F: ACGCACATCTTCCTTCGCTA, R: GCAAAAGGATCGGACACTGC; product 700 bp [54]) and female marker Lp_0170_3 (F: CAGCATTCTTGACGCACAGA, R: CCAGTGCCCCAAACAAGTAC; product 450 bp; this study). Sex-specific nucleotide sequences were identified by alignment of ancestral sex-linked gene sequences of *Ectocarpus* [55,56] to the *L. pallida* genome. PCR primers were then designed using Primer3.

PCR was performed using KAPA3G polymerase with both markers present in each reaction, on DNA extracted from two independent sets of female and male wild type gametophytes (control), the gametophyte strains tested in this study, and crossed and parthenogenetic sporophyte tissue. In the PCR, initial denaturation lasted for 3 min at 95°C, followed by 35 cycles of denaturation at 95°C for 20 sec, annealing at 61°C for 25 sec (first ten cycles touchdown 66 to 61°C), elongation at 72°C for 15 sec, and a final elongation step at 72°C for 5 min. Sex marker amplification confirmed the genetic sex of the cultures and the presence of only the male marker in male sporophytes (**Figure S1C**).

### Genome sequencing, assembly and annotation

Genomic HMW DNA was isolated from *L. pallida* strains NAM93 female and KU1234 female (**Supplementary Table S1**) using the OmniPrep Genomic DNA Isolation kit (G-Biosciences, St. Louis, USA) following the manufacturer’s instructions. Sequencing libraries were prepared using Oxford Nanopore’s SQK-LSK110 kit and sequenced on a R9.4.1 MinION flowcell (FLO-MIN106). The initial basecalling was performed using Guppy v6.0.7+c7819bc52 with the r9.4.1_450bps_hac configuration. A second basecalling was performed by Dorado v0.9.6 and the dna_r9.4.1_e8_sup@v3.6 protocol. An additional short-read sequencing library of *L. pallida* NAM93 male was prepared using a custom protocol for Tn5 tagmentation [28] and sequenced as 150 bp paired-end libraries on an Illumina NextSeq2000 at the Genome Core Facility at the MPI Tübingen Campus. A contamination analysis was performed to remove prokaryotic reads. The fastq reads were filtered for bacterial contamination by using a conjunction of Kraken2 v.2.1.2 [57] and blastn v2.9.0+ in combination with the NCBI nt database (download date 1 July 2022).

Oxford Nanopore long reads were assembled using Flye v2.9.2 [58] in default mode. Illumina short reads were individually assembled by Abyss v2.3.7 [59] (k=48 kc=3 B=50G n=5). ARCS v1.2.5 [60] (z=1500 m=8-10000 s=70 c=3 l=3 a=0.3) was used to scaffold the Nanopore Flye assembly with the Illumina Abyss assembly. Repeat soft-masking was performed by funannotate v1.8.17 [61] and default settings.

Structural gene annotation was performed by Helixer v0.3.6 [62] (--linage land_plant) and funannotate. AGAT agat_sp_merge_annotations.pl v1.5.1 [63] was used to merge the gene models of Helixer and funannotate. mRNA models spanning over >3 different gene models were removed. TE filtered protein sequences of each mRNA model were extracted and analysed by InterProScan [64], mapped by Diamond [65] to the NCBI blast nr database, Araport11 [66] and the *Ectocarpus* sp7 annotation.

### Time course of reproduction

Before the start of the experiment, gametophytes of all strains were treated with antibiotics for 14 days (Rifampicin 40 mg/L, Penicillin 100 mg/L, Streptomycin 25 mg/L, Chloramphenicol 5 mg/L, Kanamycin 50 mg/L, Cefotaxime 100 mg/L) with medium changes performed every 3-4 days. Filaments were sown to a target density of 500 gametophytes cm^−2^ and gametophyte fertility was induced as described above. Additionally, five dishes containing both the male line Lp_M1 and the female line Lp_F1 were prepared at a target density of 250 gametophytes cm^−2^ per strain to obtain crossed sporophytes. Per strain, five replicate dishes were prepared (n=5).

Development was assessed by microscopic counting (Axio Vert.A1, Zeiss, Jena, Germany) of least 300 individuals per replicate on days 0, 6, 15, 27, and 40 of the experiment, and assigning them to developmental stages (male: vegetative, mature, transition towards parthenogenesis; female: vegetative, oogonia, egg release, gametophyte with attached sporophyte). Additionally, the total number of emerging (partheno-)sporophytes was counted. Male gametophytes were categorized as transition phenotypes if they began to produce enlarged cells resembling oogonia, released enlarged immotile gametes, and/or produced sporophyte tissue from a gametophyte colony (see “Transition” category in **Figure 1B,D**). Biomass was collected at five developmental stages on days 1, 7, 16, 28, and 41: 1. Vegetative gametophytes; 2. Mature gametophytes with gametangia and first spontaneous gamete release; 3. Gametophytes after gamete release initiating vegetative growth; 4. Gametophytes during phenotype transition; in the male cultures this corresponds to reproductive variations as explained above, in the female wild type and crossing culture this corresponds to 2-5 cell (partheno-)sporophytes; 5. Blades of parthenogenetic and crossed sporophytes after syngamy (**Figure 1A,B**). Per replicate dish, 5-10 gametophyte filaments were collected using a drawn-out Pasteur pipette. Isolates were transferred in a 1 µL droplet of SW into 5 µL of lysis buffer (NEBNext® Single Cell/Low Input RNA Library Prep Kit for Illumina), and processed according to the manufacturer’s instructions. A total of 99 cDNA libraries (*n* = 3) were sequenced on an Illumina NextSeq2000 as 150 bp paired-end reads.

### Gene expression analysis

Reads from 99 low-input RNAseq libraries were mapped to the *L. pallida* genome using HiSat2 v2.2.1 [67] (--very-sensitive --new-summary -q --max-intronlen 25000). Gene read abundances were summarised using featureCounts v2.1.1 [68] (-p --countReadPairs -C -s 0 -M -O --largestOverlap –primary). Transcript abundances were calculated as transcript per million (TPM) and normalized as log2(TPM+1). Using a per-library expression threshold of TPM values > 5th percentile, excluding genes with null expression, we identified all 39955 mRNA transcripts as expressed in at least one library. Therefore, all mRNA transcripts were included in the downstream analyses.

Differential gene expression was analysed using DESeq2 v1.46.0 [69]. The significance threshold for differentially expressed genes was considered at log2 fold-change |log2FC| > 1 and FDR-adjusted padj < 0.001. Sex-biased genes were defined for each strain as the set of DEGs between the respective male and the female gametophyte Lp_F1 during maturity. For Lp_F1, Lp_M1 was chosen as the male comparison. Differences in normalised expression between female- and male-biased genes along development were analysed using pairwise Wilcoxon rank sum tests.

A weighted gene co-expression network analysis was conducted using R package WGCNA v1.73 [70] using log_2_(TPM+1) normalized data as input. Genes with null expression were excluded from the analysis, and one library was excluded as an outlier due to lack of sample clustering, leaving 98 low-input RNAseq libraries for the analysis. A signed network was constructed using a biweight midcorrelation (‘‘bicor’’), a soft thresholding power of 9, and maximum portion of outliers of 0.05. To obtain a reasonable number of gene co-expression modules with sufficient size for statistical analyses, modules were constructed with a minimum size of 30 transcripts and were merged to a correlation threshold of 0.75. We correlated gene co-expression module eigengenes to traits related to life cycle stage (gametophyte, sporophyte), genetic sex (female, male), and developmental stages for each gametophyte sex and zygotic embryo development (**see Figure 3C**). Significant enrichment of gene ontology (GO) terms within co-expression modules and differentially expressed gene sets was analysed using Fisher’s exact test (p < 0.05) within R package topGO version 2.58.0 [71].

### Field sample collection

To identify potential signals of clonality in field populations of sporophytes, we collected sporophyte tissue from *L. pallida* populations close to Swakopmund, Namibia (NAM, −22.699571 N, 14.520699 E) and Paternoster, South Africa (ZAF, −32.81789 N, 17.85747 E; and −32.81288 N, 17.86865 E; **Figure S2A**) in February and March 2022. At each location, we applied a hierarchical sampling design considering areas separated by three spatial levels: hundreds of metres (∼300 m), tens of metres (∼20 m), and few metres (∼5 m), in which we collected samples from at least 20 individuals in each of eight areas (∼2 m²) within the same population (**Figure S2B**). When increasing the sampled area to ensure a sufficient number of samples (up to approx. 25 m²), distances between areas were increased, too, to maintain a hierarchical design. In NAM, four areas were adjusted to transects of 15–30 m length to accommodate for the shore geography and population patchiness. In ZAF, 393 sporophytes were sampled, and 180 in NAM. Using cork borers, tissue discs of approx. 20 mm diameter were cut above the meristematic region of adult individuals whereas smaller individuals (< 15 cm) were removed whole. Samples were collected in ziplock bags and transported in a cool box. All tissue discs were wiped clean and rinsed in distilled water. Vegetative discs were then dried and preserved in ziplock bags filled with silica gel. Fertile sorus tissue was collected in centrifuge tubes filled with seawater and glass slides as settling substrate. Following spore release in the dark, monoclonal gametophyte isolates were produced [50] and maintained as described above.

#### DNA extraction and microsatellite amplification

DNA was extracted from dried tissue using the Nucleospin 96 Plant II kit (Macherey-Nagel, Düren, Germany) according to the manufacturer’s instructions using buffer PL1 for cell lysis. For each individual, 15 microsatellite loci were amplified to generate multi-locus genotypes. PCR reactions were carried out in 10 µL reaction volume following the protocol of Guzinski et al. [72], in which one of four fluorescent labels is added for multiplex sequencing following DNA amplification. Three to four samples were pooled, added to loading buffer including a size standard (SM594; [73]) and subsequently sequenced in an ABI 3130 XL capillary sequencer (Applied Biosystems, Waltham, USA) at Genomer, Plateforme génomique, Biological Station of Roscoff, France. Alleles were scored manually using GeneMapper v4.0 (Applied Biosystems, Waltham, USA).

Ten markers were retained for the analysis of genetic structure and diversity. Three markers had been developed for *Laminaria digitata* (CN466672 [74]; Ld19_025, Ld19_067 [75]) and seven for *Laminaria ochroleuca* (Lo5-8, LoIVVIV_15, LoIVVIV_16, LoIVVIV_24, LoIVVIV_26, LoIVVIV_27, LoIVVIV_28 [76]). Due to faulty amplification or multiple peaks, alleles of the additional markers Ld2_148, Ld19_034, Ld19_038 [75], and Ld2_357 [77] were not scored. Due to extreme polymorphism of locus LoIVVIV_23 [76] (114 alleles), this locus was removed from the analysis. Three loci showed evidence for a null allele according to the Oosterhout estimation in Micro-checker v2.2 (ZAF, Ld_19_025, frequency 0.0614; LoIVVIV_26, freq. 0.0398; NAM, LoIVVIV_24, freq. 0.0749; **Table S7**). However, as estimated null allele frequencies were low and not consistent between populations, we analysed the dataset without correction for null alleles. All markers were considered independent as no significant linkage disequilibrium could be observed within the populations (**Table S8**). The number of alleles per locus ranged between five (Lo5-8) and 31 (LoIVVIV_24). Only samples in which at least 8 markers amplified were retained for analysis.

#### Genetic diversity

The presence of null alleles was tested using the software Micro-checker v2.2 [78]. Standard measures of genetic diversity were calculated for each of the eight areas sampled per location as number of alleles per locus (N_a_), unbiased expected heterozygosity (H_e_), observed heterozygosity (H_o_), and number of private alleles (P_a_) using GenAlEx v6.5 [79]. Significant differences of diversity indices between populations were analysed by analysis of variance (ANOVA) in R v4.2.1 [80] using package nlme [81] using microsatellite loci as replicates. Allelic richness, linkage disequilibrium between pairs of loci and estimates of deviation from random mating (F_IS_) according to [82] were calculated using FSTAT v2.9.3 [83]. F_IS_ was computed within each level of sampling hierarchy [84] for all populations, and for the populations of ZAF and NAM separately. We also report single locus F_IS_ values for ZAF and NAM as suggested by [85,86] since the distribution of their values can be a signature of clonality. Significance of linkage disequilibrium was assessed using 1800 permutations, and significant deviation of F_IS_ from 0 was assessed using a t-test (n = 10 loci). The standardised index of association between loci (rld; [87,88]) was computed using Poppr v2.9.7 [89] and 999 permutations were applied to test whether loci were randomly associated. The distribution of clonal membership (Pareto ß [90]) could not be computed due to lack of repeated multilocus genotypes.

#### Population structure

Genetic substructure was analysed using several methods. Genetic differentiation between ZAF and NAM was calculated as pairwise fixation index F_ST_ [82] using FSTAT v2.9.3 with 20 permutations (F_ST_ between quadrats with 5520 permutations). Structure v2.3.4 [91] was run with 20 iterations, a burn-in period of 10^5^ and a Markov chain Monte Carlo of 10^6^. Structure Harvester Web v0.6.94 was used to identify the most likely number of clusters K [92]. Structure analysis was repeated on the identified subpopulations to identify potential hierarchical structure. Replicates of Structure runs were combined using CLUMPP v1.1.2 [93] and bar plots were created with Distruct v1.1 [94], using CLUMPAK Web v1.1 [95].

### Mate exclusion experiment

To assess whether mate presence has an effect on asexual reproduction of gametophytes, we prepared 90 mm petri dishes by fixing a fitted 5 µm nylon filter (Spectrum Labs Spectra/Mesh) across the diameter using food-grade duplication silicone (Wagnersil 32N, Wagner Dental GmbH & Co. KG, Hückelhoven, Germany). This allowed us to follow male gametophyte development in the presence of females without confounding the observations by presence of sporophytes or by false positive categorisations of female gametophytes as transitioning males. To this end, we seeded five replicate dishes of each male strain Lp_M1 – Lp_M4, and the female wild type Lp_F1 on either side of the mesh as a negative control (absence of mate) using the method described above. As a mate presence treatment, we seeded further five replicate dishes of the four male strains only on one side, while the other side was seeded with the female wild type.

After a two-day period of recovery and settlement, dishes were placed on a shaker (Heidolph Promax 2020) at 50 rpm to ensure the medium traversed the filter (tested in advance using a dye, medium was mixed in < 1 h). Apart from this, the experiment was conducted as above and development of male gametophytes was assessed in the presence and absence of females. In the dishes with Lp_M1 and Lp_F1 both males and females were assessed to also investigate the effects of male mate presence on female development.

### Statistical analysis

Developmental count data from the time course and mate exclusion experiments were analysed in R v4.2.1 [80] using linear mixed effects models (package nlme v3.1-167 [81]) and post-hoc tests were conducted using emmeans v1.10.7 [96]. Data of male transition phenotypes across strains and time were subjected to the variance-stabilising square-root transformation to account for variance heterogeneity in the model residuals [97]. To reduce inflation with zero-values in the linear models, the first time point where the respective phenotype (transition, sporophyte, egg release) occurred significantly was determined by a t-test. The models were then fitted on all data obtained from this time point onwards.

#### Time course of reproduction

Percentages of male transition phenotypes were modelled in response to the fixed interactive factors strain and time with random slopes assigned to each replicate to account for the repeated measurement design. Fractions of sporophytes per gametophyte were modelled for each strain and cross in response to time, with random slopes assigned to each replicate. Final number of sporophytes per gametophyte on day 40 was modelled in response to strain.

#### Mate exclusion experiment

Percentages of male transition phenotypes and female gametophytes with released eggs were for each strain modelled in response to the fixed interactive factors mate presence and time with random slopes assigned to each replicate to account for the repeated measurement design. Fraction of sporophytes per gametophyte were modelled for each strain and cross in response to the interaction of partner presence and time, with random slopes assigned to each replicate. The number of female eggs released per gametophyte in response to partner presence was compared using a t-test.

## Acknowledgements

We thank Gareth Pearson and the students of the University of Namibia, Sam Nujoma Campus, Henties Bay, for assistance in field sampling, Philippe Potin, Lucie Jaugeon and Jérôme Coudret for logistical support at the Roscoff Biological Station, and Martin Gachenot for support in the ploidy analysis. We thank Agnes Henschen, Dorothee Koch and Andrea Mireille Belkacemi for support in culture maintenance. We thank the Kobe University Macroalgal Culture Collection and Inka Bartsch for sharing *L. pallida* strains. Wild samples were collected with local authorisation under material transfer agreements, following the Nagoya protocol on access and benefit sharing (South Africa, ABSCH-IRCC-ZA-264230-1; Namibia, Research Permit RPIV02022022 and Access Permit ABS-0061CHM1022).

## Data availability

The RNAseq and DNAseq data generated for this study have been deposited at NCBI (BioProject ID PRJNA1479194). The *L. pallida* genome and annotation files, count data of developmental stages over time, sporophyte areas over time, microsatellite genotype data, and raw gel image files have been deposited at EDMOND (xxxxx).

## Author contributions

DL: Investigation (lead); Formal analysis (lead); Conceptualization (supporting); Funding acquisition (supporting); Methodology (equal); Writing – original draft (lead), Writing – review and editing (supporting).

RL, LK, JJB, MDR: Investigation (equal); Methodology (supporting)

MV: Conceptualization (supporting); Methodology (equal)

FBH: Data curation (lead)

SMC: Conceptualization (lead); Funding acquisition (lead); Methodology (equal); Project administration (lead); Writing – original draft (supporting); Writing – review and editing (lead).

## Supplemental information

**Figure S1.**
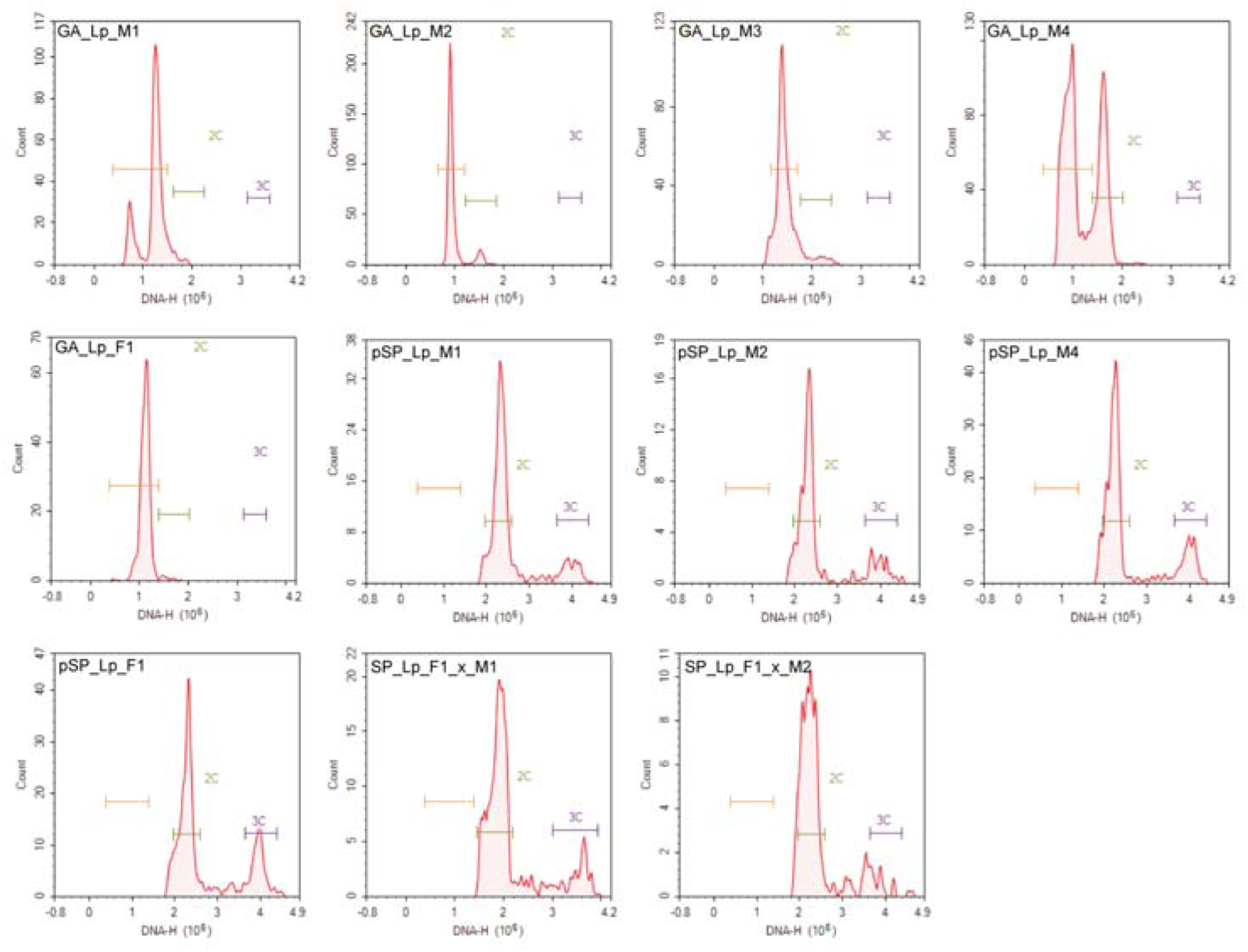
Results of flow cytometry analysis to estimate DNA content of *L. pallida* gametophyte (GA), partheno-sporophyte (pSP) and sporophyte (SP) material of five strains (males M1, M2, M3, M4; female F1; cross F1_x_M1, F1_x_M2). Plots show density of fluorescence signals (i.e., number of nuclei) as a function of DNA content (DNA-H).

**Figure S2.**
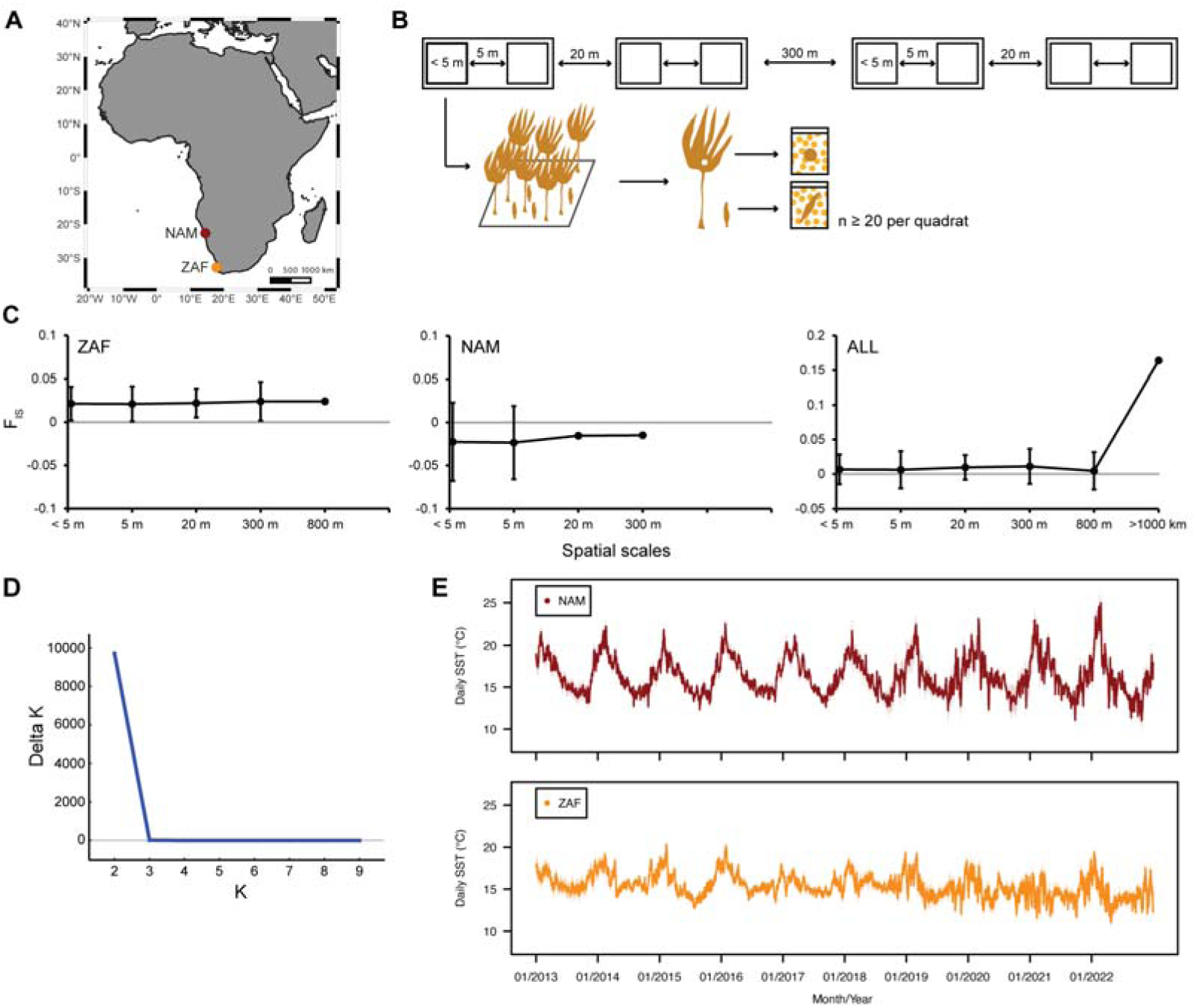
Wild populations of *Laminaria pallida* investigated in this study. **(A)** Map depicting the location of Swakopmund, Namibia (NAM) and Paternoster, South Africa (ZAF). **(B)** Schematic hierarchical sampling design applied at the two populations. **(C)** Inbreeding coefficient over spatial scales in the hierarchical sampling design for ZAF, NAM, and both populations (mean ± 95% CI). **(D)** ΔK (Evanno et al., 2005) plotted against number of genetic clusters (K), for K = 2 to K = 9 obtained with Structure Harvester. **(E)** Variation of daily sea surface temperatures for NAM and ZAF from 2013 to 2022 (10 years before time of sampling) obtained from the E.U. Copernicus Marine Service (https://doi.org/10.48670/moi-00165) with daily means shown as a solid line and shading indicating standard error.

**Figure S3.**
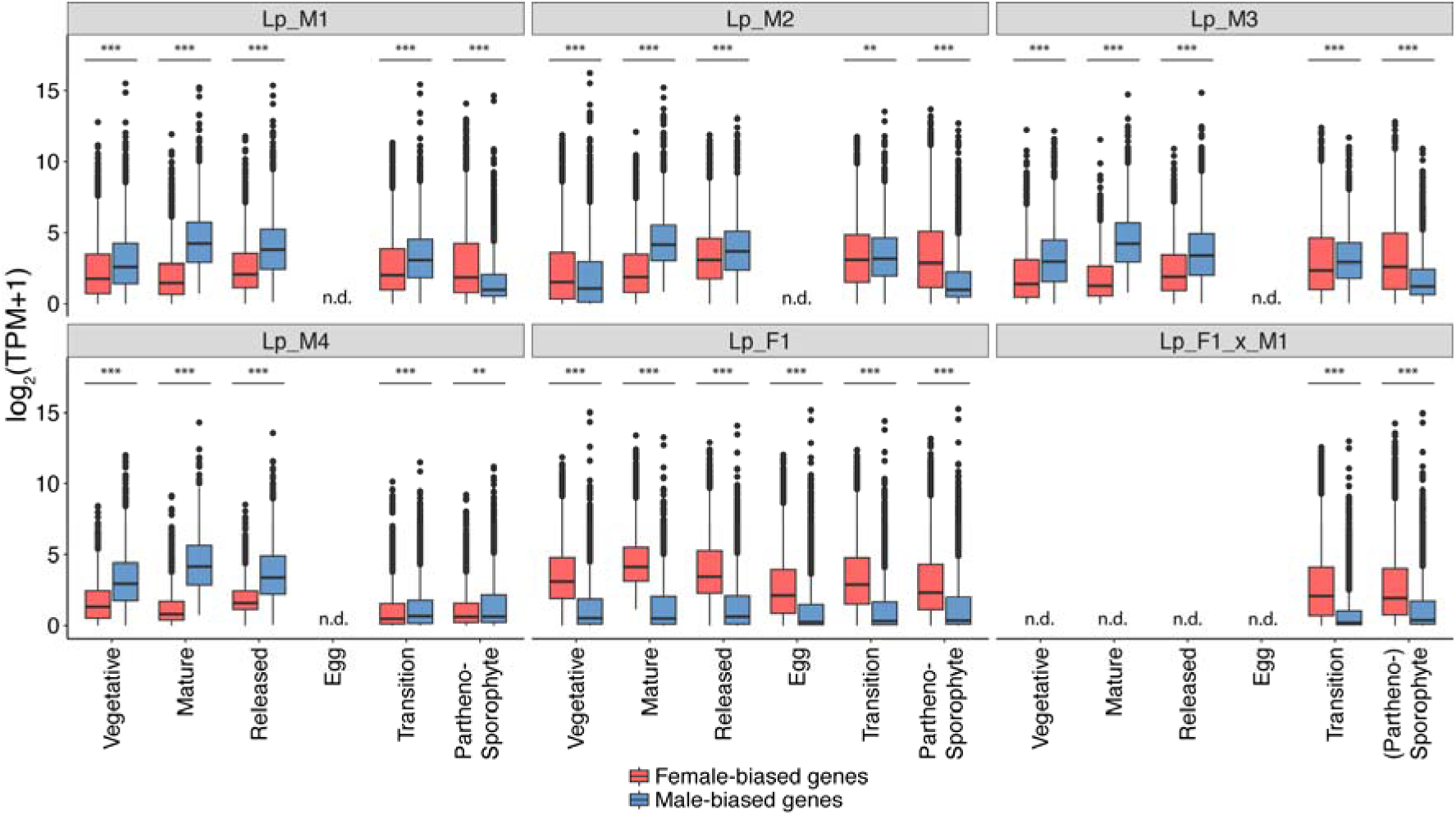
Expression of sex-biased genes along development for four male (M1, M2, M3, M4) and one female strain (F1) of *Laminaria pallida* and their cross (F1_x_M1). Significant differences between male- and female-biased gene expression are indicated with asterisks (Wilcoxon rank sum tests; **, p<0.01; ***, p<0.001); n.d., no data.

**Figure S4.**
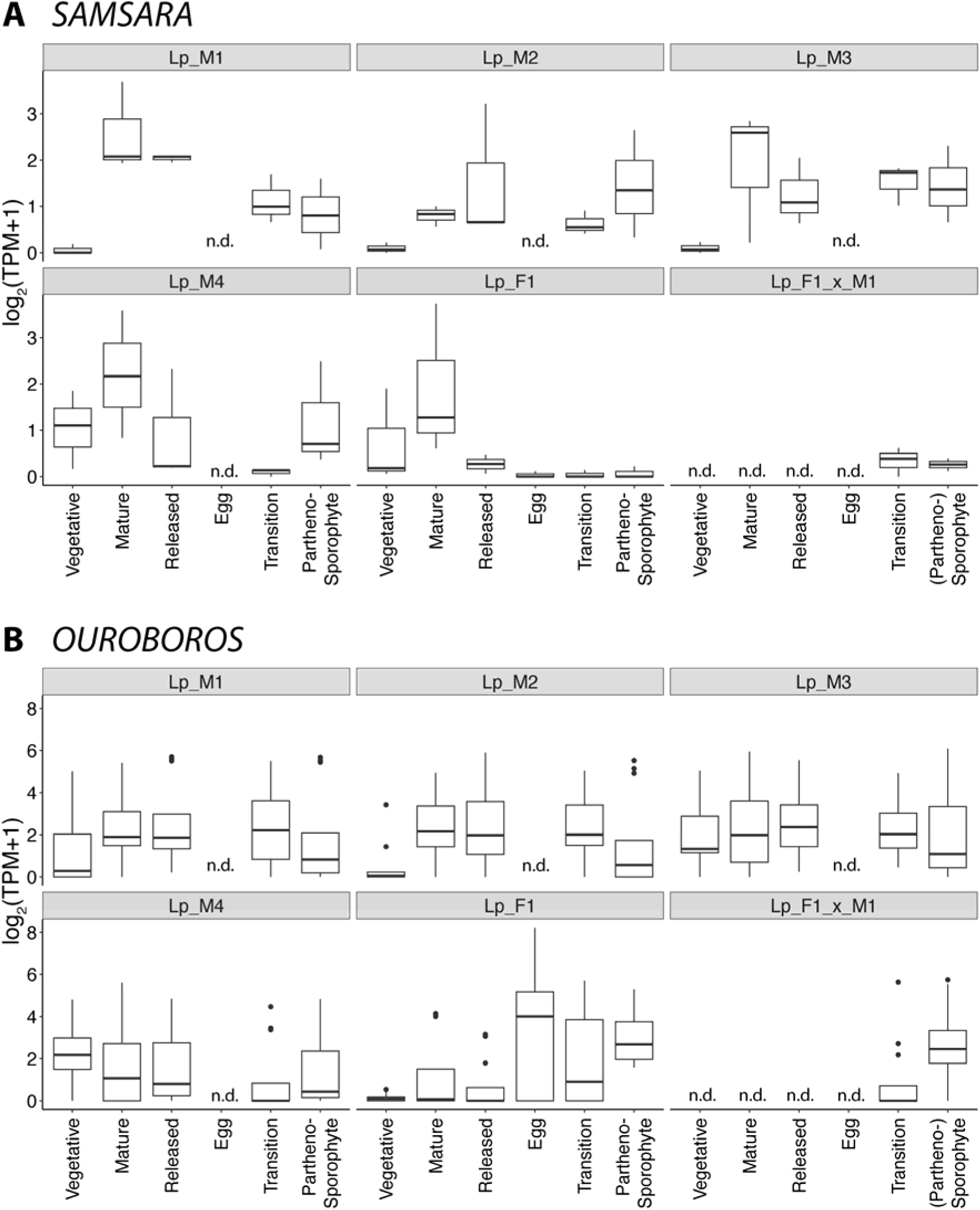
Expression of life-cycle related TALE homeodomain transcription factors (A) SAMSARA (1 transcript in n=3 replicates) and (B) OUROBOROS (4 transcripts in n=3 replicates) along development for four male (M1, M2, M3, M4) and one female strain (F1) of *Laminaria pallida* and their cross (F1_x_M1); n.d., no data.

**Figure S5.**
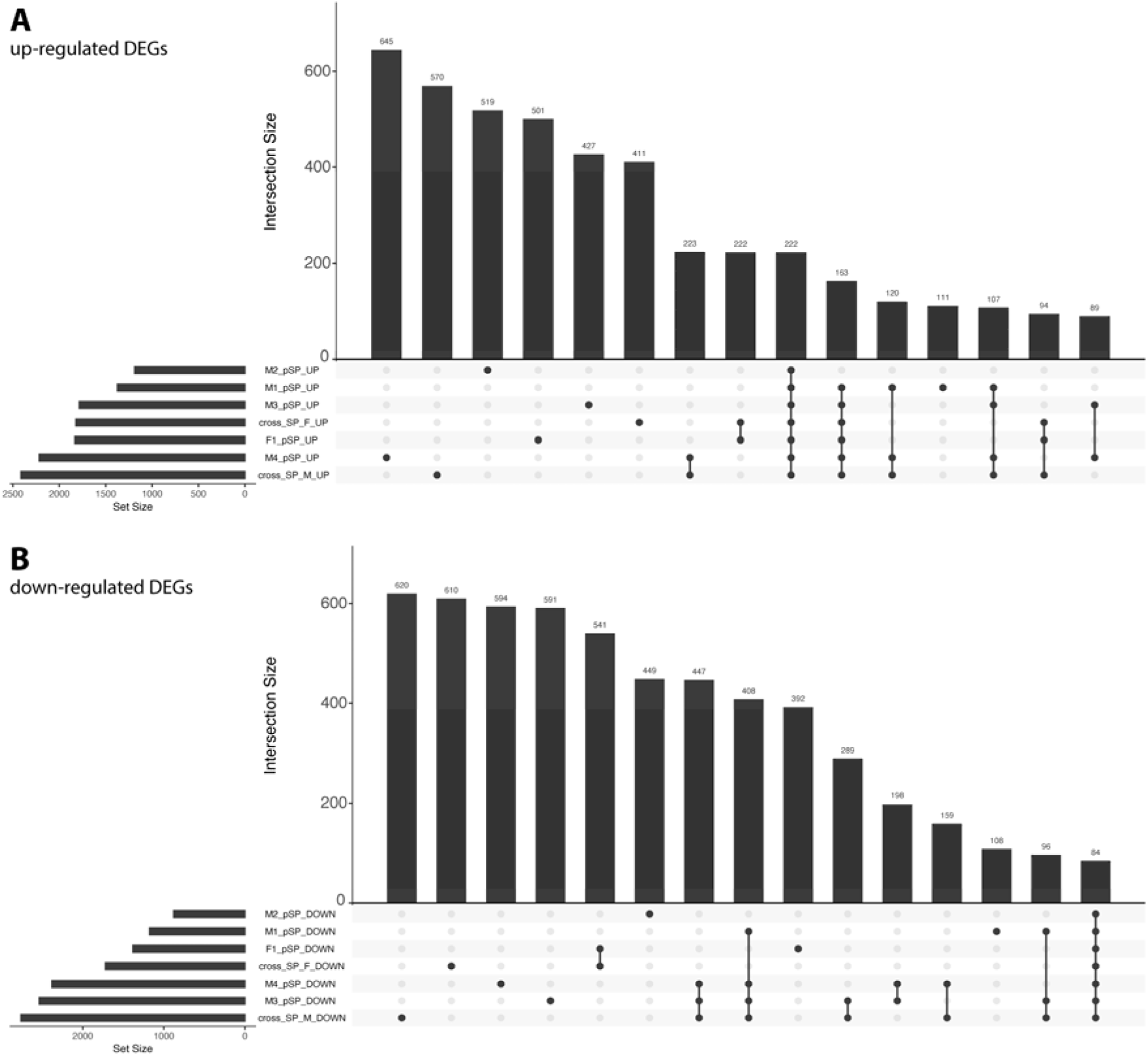
Intersection of **(A)** up-regulated and **(B)** down-regulated differentially expressed genes (DEGs) between (partheno-)sporophytes and their corresponding gametophyte for four male (M1, M2, M3, M4) and one female strain (F1) of *Laminaria pallida* and their cross (F1_x_M1). Only the 15 largest intersections are shown.

**Figure S6.**
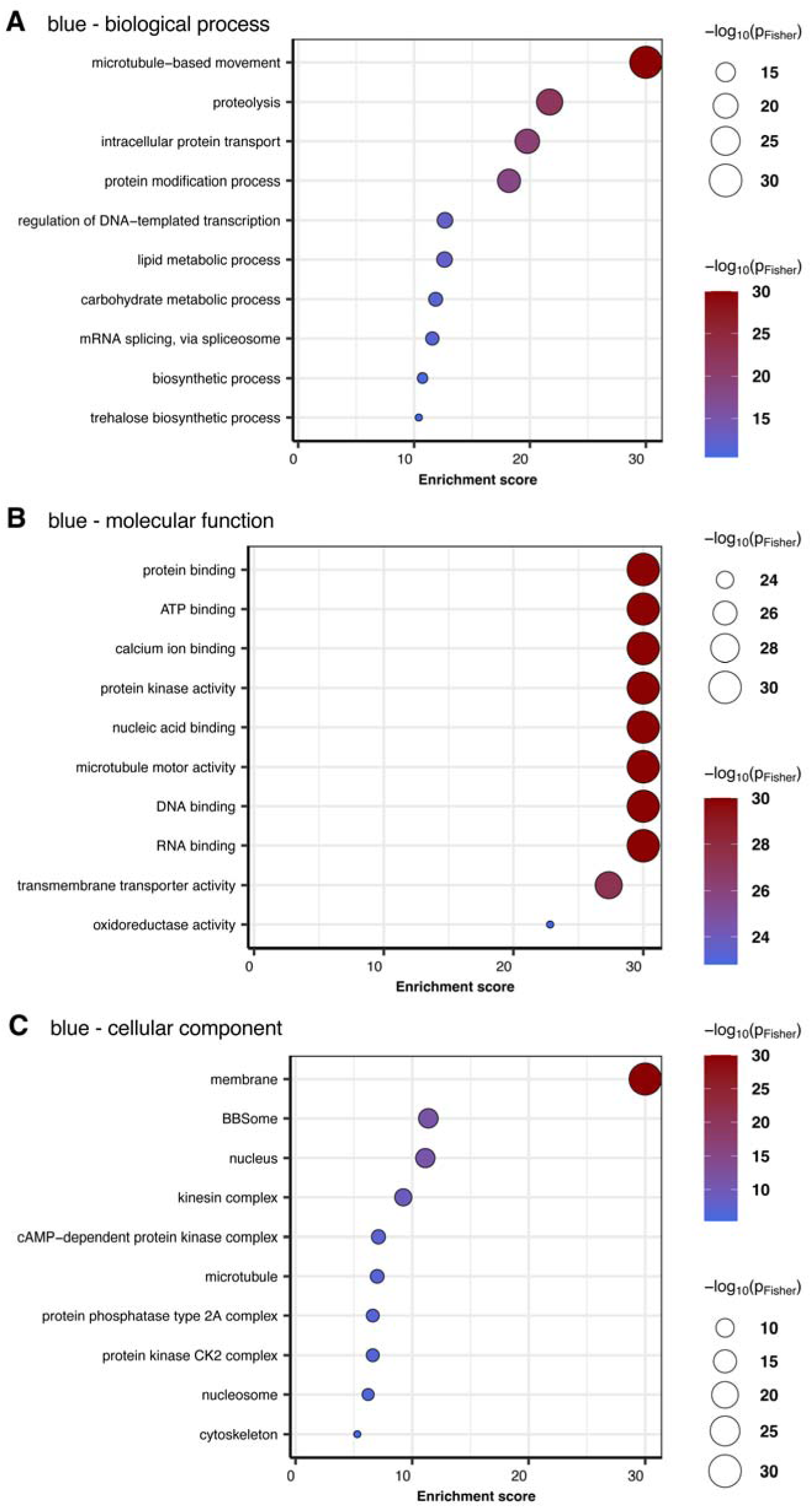
Enriched gene ontology (GO) terms for **(A)** biological process, **(B)** molecular function, **(C)** cellular component in module “blue” of weighted gene co-expression network analysis of *Laminaria pallida* development. A maximum of ten most enriched terms is displayed in dotplots sorted by significance (Fisher’s exact test).

**Figure S7.**
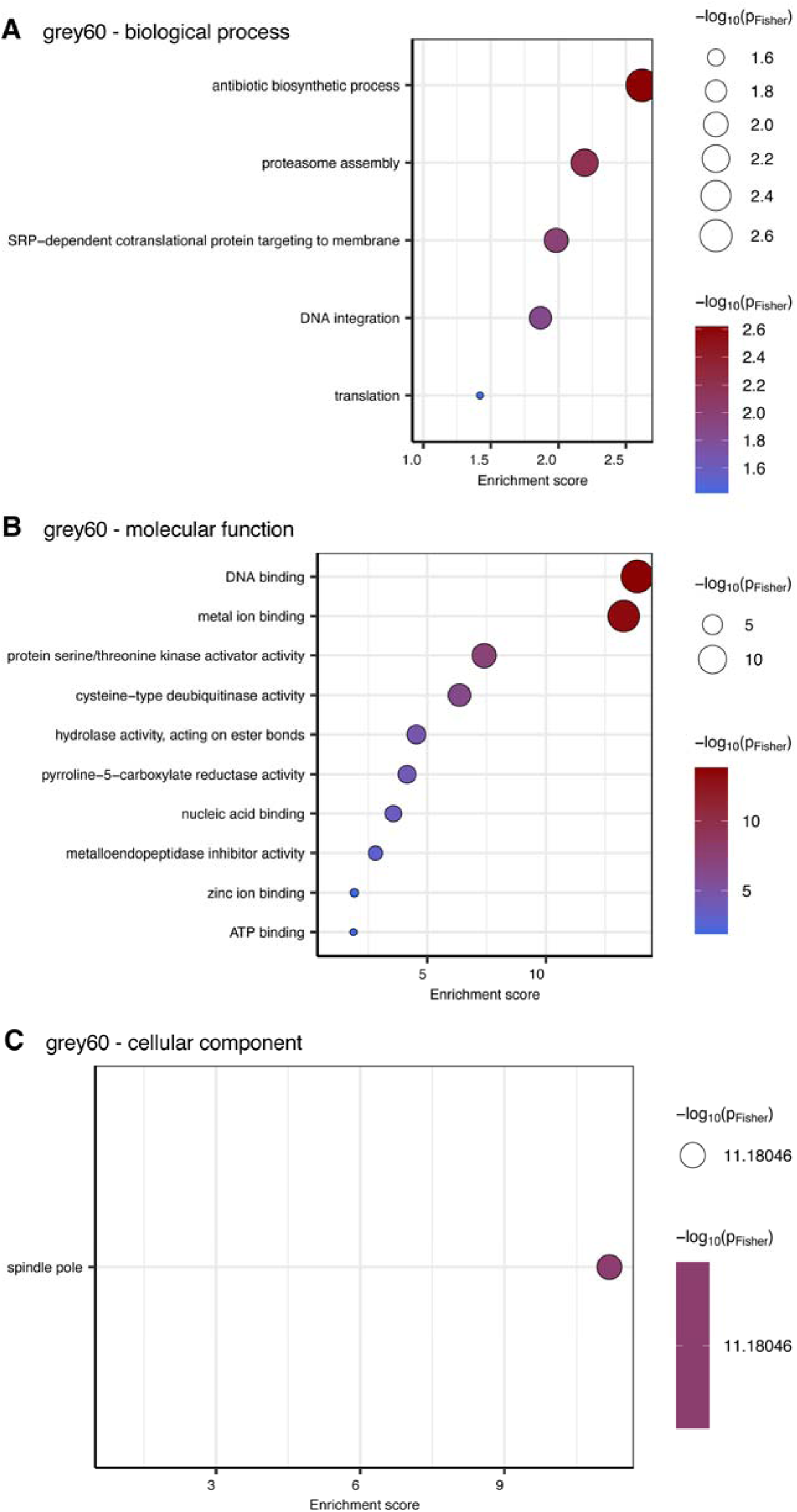
Enriched gene ontology (GO) terms for **(A)** biological process, **(B)** molecular function, **(C)** cellular component in module “grey60” of weighted gene co-expression network analysis of *Laminaria pallida* development. A maximum of ten most enriched terms is displayed in dotplots sorted by significance (Fisher’s exact test).

**Figure S8.**
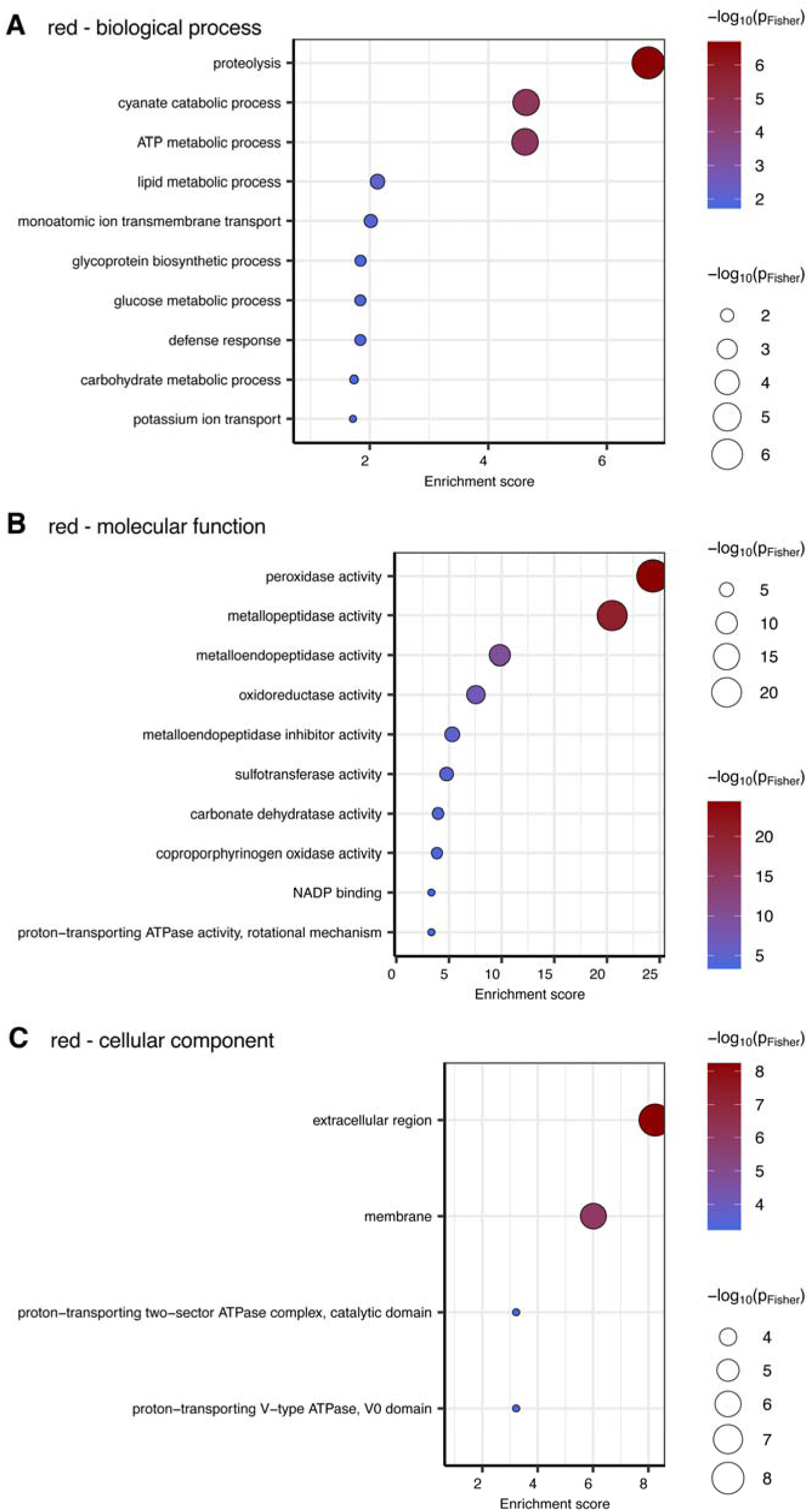
Enriched gene ontology (GO) terms for **(A)** biological process, **(B)** molecular function, **(C)** cellular component in module “red” of weighted gene co-expression network analysis of *Laminaria pallida* development. A maximum of ten most enriched terms is displayed in dotplots sorted by significance (Fisher’s exact test).

**Figure S9.**
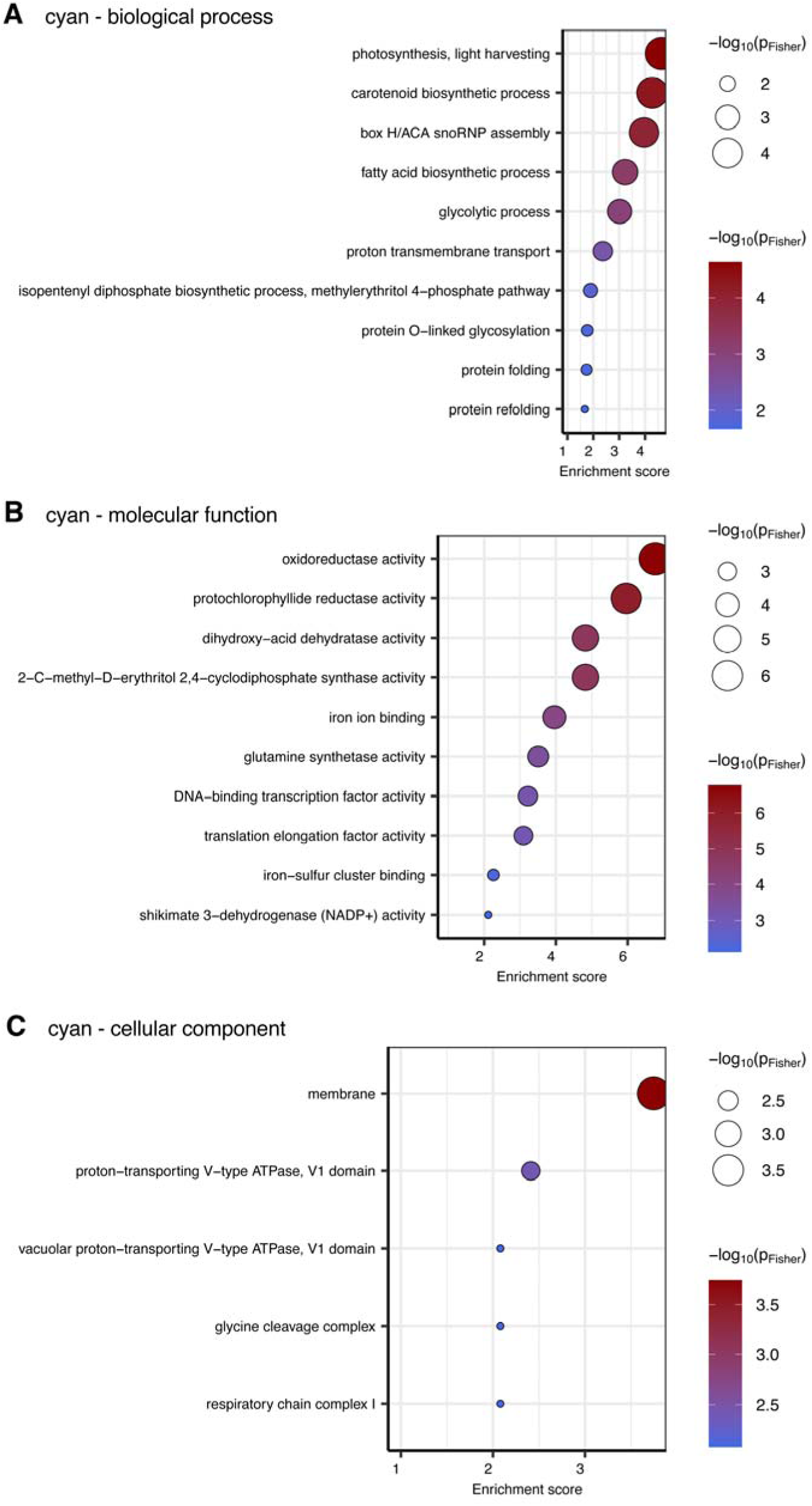
Enriched gene ontology (GO) terms for **(A)** biological process, **(B)** molecular function, **(C)** cellular component in module “cyan” of weighted gene co-expression network analysis of *Laminaria pallida* development. A maximum of ten most enriched terms is displayed in dotplots sorted by significance (Fisher’s exact test).

**Table S1.** Description of *L. pallida* gametophyte lines used in the experiment.

**Table S2.** Results of linear mixed effects models on responses during the developmental time course of *Laminaria pallida*.

**Table S3.** Expression data and functional annotation of all transcripts of *Laminaria pallida*. Functional annotation based on blastx against the *Ectocarpus* Ec32 genome, nr and Araport11 databases, gene ontology (GO) terms, normalized read counts (transcripts per million, TPM) strains Lp_M1, Lp_M2, Lp_M3, Lp_M4, Lp_F1 and cross Lp_F1_x_M1, along developmental stages vegetative (veg), mature (gam), released (rel), transition (tra) and sporophyte blade (bla), shared sporophyte-related genes based on DESeq2 results (sporophyte-gametophyte comparison), gene affiliation to gene co-expression modules, assignment to gene co-expression modules based on weighted gene co-expression analysis (WGCNA).

**Table S4.** Exact locations of the *L. pallida* sampling sites.

**Table S5.** Obervations of parthenogenesis during prolonged cultivation of unialgal gametophyte cultures isolated from *L. pallida* field material.

**Table S6.** Results of linear mixed effects models on responses during the mate exclusion experiment on *Laminaria pallida* gametophytes.

**Table S7.** Null allele estimation (Oosterhout estimate within micro-checker) for each microsatellite locus for populations of *Laminaria pallida* from Swakopmund, Namibia (NAM) and Paternoster, South Africa (ZAF). Significant estimates are highlighted in bold font.

**Table S8.** Linkage disequilibrium estimation within FSTAT for each pair of microsatellite loci for populations of *Laminaria pallida* from Swakopmund, Namibia (NAM), Paternoster, South Africa (ZAF), and both populations. D’ values and p-values are given. Adjusted p-value based on 1800 permutations is 0.00056.

## References

1. Otto SP. The evolutionary enigma of sex. Am Nat. 2009;174: S1–S14. doi:10.1086/599084

2. Williams GC. Sex and evolution. Princeton University Press; 1975.

3. Maynard Smith J. The evolution of sex. Cambridge University Press; 1978.

4. Burt A. Perspective: sex, recombination, and the efficacy of selection-Was Weismann right? Evolution. 2000;54: 337–351. doi:10.1111/j.0014-3820.2000.tb00038.x

5. Charlesworth B. The evolutionary biology of sex. Curr Biol. 2006;16: R693–R695. doi:10.1016/j.cub.2006.08.023

6. Simon J-C, Delmotte F, Rispe C, Crease T. Phylogenetic relationships between parthenogens and their sexual relatives: the possible routes to parthenogenesis in animals: Routes to parthenogenesis in animals. Biol J Linn Soc. 2003;79: 151–163. doi:10.1046/j.1095-8312.2003.00175.x

7. Hörandl E. Apomixis and the paradox of sex in plants. Ann Bot. 2024;134: 1–18. doi:10.1093/aob/mcae044

8. Barrett SCH. The evolution of plant reproductive systems: how often are transitions irreversible? Proc R Soc B. 2013;280: 20130913. doi:10.1098/rspb.2013.0913

9. Tilquin A, Kokko H. What does the geography of parthenogenesis teach us about sex? Phil Trans R Soc B. 2016;371: 20150538. doi:10.1098/rstb.2015.0538

10. Hörandl E. Geographical parthenogenesis: opportunities for asexuality. In: Schön I, Martens K, Dijk P, editors. Lost Sex: The Evolutionary Biology of Parthenogenesis. Dordrecht: Springer Netherlands; 2009. pp. 161–186. doi:10.1007/978-90-481-2770-2_8

11. Mignerot L, Avia K, Luthringer R, Lipinska AP, Peters AF, Cock JM, et al. A key role for sex chromosomes in the regulation of parthenogenesis in the brown alga *Ectocarpus*. PLoS Genet. 2019;15: e1008211. doi:10.1371/journal.pgen.1008211

12. Cossard GG, Godfroy O, Nehr Z, Cruaud C, Cock JM, Lipinska AP, et al. Selection drives convergent gene expression changes during transitions to co-sexuality in haploid sexual systems. Nat Ecol Evol. 2022;6: 579–589. doi:10.1038/s41559-022-01692-4

13. Barrera-Redondo J, Lipinska AP, Liu P, Dinatale E, Cossard G, Bogaert K, et al. Origin and evolutionary trajectories of brown algal sex chromosomes. Nat Ecol Evol. 2025. doi:10.1038/s41559-025-02838-w

14. Heesch S, SerranoLJSerrano M, BarreraLJRedondo J, Luthringer R, Peters AF, Destombe C, et al. Evolution of life cycles and reproductive traits: Insights from the brown algae. J Evol Biol. 2021;34: 992–1009. doi:10.1111/jeb.13880

15. Hoshino M, Cossard G, Haas FB, Kane EI, Kogame K, Jomori T, et al. Parallel loss of sexual reproduction in field populations of a brown alga sheds light on the mechanisms underlying the emergence of asexuality. Nat Ecol Evol. 2024;8: 1916–1932. doi:10.1038/s41559-024-02490-w

16. Coelho SM, Godfroy O, Arun A, Le Corguillé G, Peters AF, Cock JM. *OUROBOROS* is a master regulator of the gametophyte to sporophyte life cycle transition in the brown alga *Ectocarpus*. Proc Natl Acad Sci USA. 2011;108: 11518–11523. doi:10.1073/pnas.1102274108

17. Arun A, Coelho SM, Peters AF, Bourdareau S, Pérès L, Scornet D, et al. Convergent recruitment of TALE homeodomain life cycle regulators to direct sporophyte development in land plants and brown algae. eLife. 2019;8: e43101. doi:10.7554/eLife.43101

18. Teagle H, Hawkins SJ, Moore PJ, Smale DA. The role of kelp species as biogenic habitat formers in coastal marine ecosystems. J Exp Mar Biol Ecol. 2017;492: 81–98. doi:10.1016/j.jembe.2017.01.017

19. tom Dieck (Bartsch) I. North Pacific and North Atlantic digitate *Laminaria* species (Phaeophyta): hybridization experiments and temperature responses. Phycologia. 1992;31: 147–163. doi:10.2216/i0031-8884-31-2-147.1

20. Martins N, Tanttu H, Pearson GA, Serrão EA, Bartsch I. Interactions of daylength, temperature and nutrients affect thresholds for life stage transitions in the kelp *Laminaria digitata* (Phaeophyceae). Bot Mar. 2017;60. doi:10.1515/bot-2016-0094

21. Dries E, Meyers Y, Liesner D, Gonzaga FM, Becker JFM, Zakka EE, et al. Cell wallLJmediated maternal control of apical–basal patterning of the kelp *Undaria pinnatifida*. New Phytol. 2024;243: 1887–1898. doi:10.1111/nph.19953

22. Nakahara H, Nakamura Y. Parthenogenesis, apogamy and apospory in *Alaria crassifolia* (Laminariales). Mar Biol. 1973;18: 327–332. doi:10.1007/BF00347797

23. Ar Gall E, Asensi A, Marie D, Kloareg B. Parthenogenesis and apospory in the Laminariales: A flow cytometry analysis. Eur J Phycol. 1996;31: 369–380. doi:10.1080/09670269600651601

24. Müller DG, Murúa P, Westermeier R. Reproductive strategies of *Lessonia berteroana* (Laminariales, Phaeophyceae) gametophytes from Chile: apogamy, parthenogenesis and cross-fertility with *L. spicata*. J Appl Phycol. 2019;31: 1475–1481. doi:10.1007/s10811-018-1625-9

25. Druehl LD, Collins JD, Lane CE, Saunders GW. An evaluation of methods used to assess intergeneric hybridization in kelp using Pacific Laminariales (Phaeophyceae). J Phycol. 2005;41: 250–262. doi:10.1111/j.1529-8817.2005.04143.x

26. Destombe C, Oppliger LV. Male gametophyte fragmentation in *Laminaria digitata*: a life history strategy to enhance reproductive success. Cah Biol Mar. 2011;52: 385–394.

27. Liboureau P, Pearson GA, Serrão EA, Kreiner A, Martins N. A novel sexual system in male gametophytes of *Laminaria pallida* (Phaeophyceae). European Journal of Phycology. 2024;59: 232–241. doi:10.1080/09670262.2024.2314487

28. Liesner D, Cossard GG, Zheng M, Godfroy O, Barrera-Redondo J, Haas FB, et al. Developmental pathways underlying sexual differentiation in the U/V sex chromosome system of giant kelp. Dev Cell. 2025;60: 1142–1152.e6. doi:10.1016/j.devcel.2024.12.022

29. Evanno G, Regnaut S, Goudet J. Detecting the number of clusters of individuals using the software structure: a simulation study. Mol Ecol. 2005;14: 2611–2620. doi:10.1111/j.1365-294X.2005.02553.x

30. Oppliger LV, Correa JA, Peters AF. Parthenogenesis in the brown alga *Lessonia nigrescens* (Laminariales, Phaeophyceae) from central Chile. J Phycol. 2007;43: 1295–1301. doi:10.1111/j.1529-8817.2007.00408.x

31. Yue S, Xue N, Yi C, Sun J, Li X, Chen S, et al. Apomixis in *Saccharina japonica*: parthenogenesis in male and apogamy in female gametophytes. Aquaculture. 2024;591: 741142. doi:10.1016/j.aquaculture.2024.741142

32. Zhang L, Xue N, Li X, Zhou X, Yang G. Apomixis in kelp genetic improvement: Practices, challenges, and prospects. Aquaculture. 2025;598: 741996. doi:10.1016/j.aquaculture.2024.741996

33. Tatarenkov A, Bergström L, Jönsson RB, Serrão EA, Kautsky L, Johannesson K. Intriguing asexual life in marginal populations of the brown seaweed *Fucus vesiculosus*. Mol Ecol. 2005;14: 647–651. doi:10.1111/j.1365-294X.2005.02425.x

34. Pereyra RT, Kinnby A, Le Moan A, OrtegaLJMartinez O, Jonsson PR, Piarulli S, et al. An evolutionary mosaic challenges traditional monitoring of a foundation species in a coastal environment — the Baltic *Fucus vesiculosus*. Mol Ecol. 2025; e17699. doi:10.1111/mec.17699

35. Demes KW, Graham MH. Abiotic regulation of investment in sexual versus vegetative reproduction in the clonal kelp *Laminaria sinclairii* (Laminariales, Phaeophyceae): ecophysiology of a clonal kelp. J Phycol. 2011;47: 463–470. doi:10.1111/j.1529-8817.2011.00981.x

36. Westermeier R, Murúa P, Patiño DJ, Muñoz L, Ruiz A, Atero C, et al. Utilization of holdfast fragments for vegetative propagation of *Macrocystis integrifolia* in Atacama, Northern Chile. J Appl Phycol. 2013;25: 639–642. doi:10.1007/s10811-012-9898-x

37. Reynes L, Thibaut T, Mauger S, Blanfuné A, Holon F, Cruaud C, et al. Genomic signatures of clonality in the deep water kelp *Laminaria rodriguezii*. Mol Ecol. 2021;30: 1806–1822. doi:10.1111/mec.15860

38. Fang Z-X, Dai J-X, Ou Y-L, Cui J-J, Chen D-Q. Some genetic observations on the monoploid breeding of *Laminaria japonica*. Scientia Sinica. 1978;21: 401–408. doi:10.1360/ya1978-21-3-401

39. Macaisne N, Liu F, Scornet D, Peters AF, Lipinska A, Perrineau M-M, et al. The *Ectocarpus IMMEDIATE UPRIGHT* gene encodes a member of a novel family of cysteine-rich proteins with an unusual distribution across the eukaryotes. Development. 2017;144: 409–418. doi:10.1242/dev.141523

40. Peters AF, Scornet D, Ratin M, Charrier B, Monnier A, Merrien Y, et al. Life-cycle-generation-specific developmental processes are modified in the *immediate upright* mutant of the brown alga *Ectocarpus siliculosus*. Development. 2008;135: 1503–1512. doi:10.1242/dev.016303

41. Coelho SMB, Brownlee C, Bothwell JHF. A tip-high, Ca^2+^-interdependent, reactive oxygen species gradient is associated with polarized growth in *Fucus serratus* zygotes. Planta. 2008;227: 1037–1046. doi:10.1007/s00425-007-0678-9

42. Ali MF, Muday GK. Reactive oxygen species are signaling molecules that modulate plant reproduction. Plant Cell Environ. 2024;47: 1592–1605. doi:10.1111/pce.14837

43. Vijverberg K, Ozias-Akins P, Schranz ME. Identifying and engineering genes for parthenogenesis in plants. Front Plant Sci. 2019;10: 128–144. doi:10.3389/fpls.2019.00128

44. Johansson ML, Raimondi PT, Reed DC, Coelho NC, Serrão EA, Alberto FA. Looking into the black box: simulating the role of selfLJfertilization and mortality in the genetic structure of *Macrocystis pyrifera*. Mol Ecol. 2013;22: 4842–4854. doi:10.1111/mec.12444

45. Mráz P, Mrázová V. Greater reproductive assurance of asexual plant compared with sexual relative in a lowLJdensity sympatric population: Experimental evidence for pollen limitation. J Evol Biol. 2021;34: 1503–1509. doi:10.1111/jeb.13910

46. Pereyra RT, Rafajlović M, De Wit P, Pinder M, Kinnby A, Töpel M, et al. Clones on the run: The genomics of a recently expanded partially clonal species. Mol Ecol. 2023;32: 4209–4223. doi:10.1111/mec.16996

47. Martins N, Pearson GA, Gouveia L, Tavares AI, Serrão EA, Bartsch I. Hybrid vigour for thermal tolerance in hybrids between the allopatric kelps *Laminaria digitata* and *L. pallida* (Laminariales, Phaeophyceae) with contrasting thermal affinities. Eur J Phycol. 2019;54: 548–561. doi:10.1080/09670262.2019.1613571

48. Oppliger LV, Von Dassow P, Bouchemousse S, Robuchon M, Valero M, Correa JA, et al. Alteration of sexual reproduction and genetic diversity in the kelp species *Laminaria digitata* at the southern limit of its range. PLoS ONE. 2014;9: e102518. doi:10.1371/journal.pone.0102518

49. Li J, Pang S. Evidence of sterility of the male sporophytes of the brown alga *Saccharina japonica* (Phaeophyceae) in culture irrespective of their ploidy levels. J Phycol. 2024; jpy.13530. doi:10.1111/jpy.13530

50. Bartsch I. Derivation of clonal stock cultures and hybridization of kelps. 1st ed. In: Charrier B, Wichard T, Reddy CRK, editors. Protocols for Macroalgae Research. 1st ed. Boca RatonLJ: Taylor & Francis, 2018.: CRC Press; 2018. pp. 61–78. doi:10.1201/b21460-3

51. Starr RC, Zeikus JA. UTEX—the culture collection of algae at the University of Texas at Austin. Journal of Phycology. 1987;23(Suppl): 1–47.

52. Tatewaki M. Formation of a crustaceous sporophyte with unilocular sporangia in *Scytosiphon lomentaria*. Phycologia. 1966;6: 62–66. doi:10.2216/i0031-8884-6-1-62.1

53. Schindelin J, Arganda-Carreras I, Frise E, Kaynig V, Longair M, Pietzsch T, et al. Fiji: an open-source platform for biological-image analysis. Nat Methods. 2012;9: 676–682. doi:10.1038/nmeth.2019

54. Luthringer R, Raphalen M, Guerra C, Colin S, Martinho C, Zheng M, et al. Repeated co-option of HMG-box genes for sex determination in brown algae and animals. Science. 2024;383: eadk5466. doi:10.1126/science.adk5466

55. Lipinska AP, Toda NRT, Heesch S, Peters AF, Cock JM, Coelho SM. Multiple gene movements into and out of haploid sex chromosomes. Genome Biol. 2017;18: 104. doi:10.1186/s13059-017-1201-7

56. Lipinska AP, Ahmed S, Peters AF, Faugeron S, Cock JM, Coelho SM. Development of PCRLJbased markers to determine the sex of kelps. PLoS ONE. 2015;10: e0140535. doi:10.1371/journal.pone.0140535

57. Wood DE, Lu J, Langmead B. Improved metagenomic analysis with Kraken 2. Genome Biol. 2019;20: 257–269. doi:10.1186/s13059-019-1891-0

58. Kolmogorov M, Yuan J, Lin Y, Pevzner PA. Assembly of long, error-prone reads using repeat graphs. Nat Biotechnol. 2019;37: 541–546. doi:10.1038/s41587-019-0072-8

59. Jackman SD, Vandervalk BP, Mohamadi H, Chu J, Yeo S, Hammond SA, et al. ABySS 2.0: resource-efficient assembly of large genomes using a bloom filter. Genome Res. 2017;27: 768–777. doi:10.1101/gr.214346.116

60. Yeo S, Coombe L, Warren RL, Chu J, Birol I. ARCS: scaffolding genome drafts with linked reads. Sahinalp C, editor. Bioinformatics. 2018;34: 725–731. doi:10.1093/bioinformatics/btx675

61. Palmer JM, Stajich J. Funannotate v1.8.1: eukaryotic genome annotation. Zenodo; 2020. doi:10.5281/zenodo.4054262

62. Holst F, Bolger AM, Kindel F, Günther C, Maß J, Triesch S, et al. Helixer: ab initio prediction of primary eukaryotic gene models combining deep learning and a hidden markov model. Nat Methods. 2025; 0–19. doi:10.1038/s41592-025-02939-1

63. Dainat J. Another Gtf/Gff Analysis Toolkit (AGAT): Resolve interoperability issues and accomplish more with your annotations. 2022. Available: https://github.com/NBISweden/AGAT

64. Jones P, Binns D, Chang H-Y, Fraser M, Li W, McAnulla C, et al. InterProScan 5: genome-scale protein function classification. Bioinform (Oxf Engl). 2014;30: 1236–1240. doi:10.1093/bioinformatics/btu031

65. Buchfink B, Reuter K, Drost H-G. Sensitive protein alignments at tree-of-life scale using DIAMOND. Nat Methods. 2021;18: 366–368. doi:10.1038/s41592-021-01101-x

66. Cheng C-Y, Krishnakumar V, Chan AP, Thibaud-Nissen F, Schobel S, Town CD. Araport11: a complete reannotation of the *Arabidopsis thaliana* reference genome. Plant J. 2017;89: 789–804. doi:10.1111/tpj.13415

67. Kim D, Paggi JM, Park C, Bennett C, Salzberg SL. Graph-based genome alignment and genotyping with HISAT2 and HISAT-genotype. Nat Biotechnol. 2019;37: 907–915. doi:10.1038/s41587-019-0201-4

68. Liao Y, Smyth GK, Shi W. featureCounts: an efficient general purpose program for assigning sequence reads to genomic features. Bioinformatics. 2014;30: 923–930. doi:10.1093/bioinformatics/btt656

69. Love MI, Huber W, Anders S. Moderated estimation of fold change and dispersion for RNA-seq data with DESeq2. Bioinformatics; 2014. doi:10.1101/002832

70. Langfelder P, Horvath S. WGCNA: an R package for weighted correlation network analysis. BMC Bioinformatics. 2008;9: 559. doi:10.1186/1471-2105-9-559

71. Alexa A, Rahnenführer J, Lengauer T. Improved scoring of functional groups from gene expression data by decorrelating GO graph structure. Bioinformatics. 2006;22: 1601–1607. doi:10.1093/bioinformatics/btl140

72. Guzinski J, Mauger S, Cock JM, Valero M. Characterization of newly developed expressed sequence tag-derived microsatellite markers revealed low genetic diversity within and low connectivity between European Saccharina latissima populations. J Appl ycol. 2016;28: 3057–3070. doi:10.1007/s10811-016-0806-7

73. Mauger S, Couceiro L, Valero M. A simple and cost-effective method to synthesize an internal size standard amenable to use with a 5-dye system. Prime Research on Biotechnology. 2012;2: 40–46.

74. Brennan G, Kregting L, Beatty GE, Cole C, Elsäßer B, Savidge G, et al. Understanding macroalgal dispersal in a complex hydrodynamic environment: a combined population genetic and physical modelling approach. J R Soc Interface. 2014;11: 20140197. doi:10.1098/rsif.2014.0197

75. Mauger S, Fouqueau L, Avia K, Reynes L, Serrao EA, Neiva J, et al. Development of tools to rapidly identify cryptic species and characterize their genetic diversity in different European kelp species. J Appl Phyco. 2021;33: 4169–4186. doi:10.1007/s10811-021-02613-x

76. Coelho NC, Serrão EA, Alberto F. Characterization of fifteen microsatellite markers for the kelp Laminaria ochroleuca and cross species amplification within the genus. Conservation Genet Resour. 2014;6: 949–950. doi:10.1007/s12686-014-0249-x

77. Billot C, Rousvoal S, Estoup A, Epplen JT, SaumitouLJLaprade P, Valero M, et al. Isolation and characterization of microsatellite markers in the nuclear genome of the brown alga *Laminaria digitata* (Phaeophyceae). Mol Ecol. 1998;7: 1778–1780. doi:10.1046/j.1365-294x.1998.00516.x

78. Van Oosterhout C, Hutchinson WF, Wills DPM, Shipley P. micro-checker: software for identifying and correcting genotyping errors in microsatellite data. Mol Ecol Notes. 2004;4: 535–538. doi:10.1111/j.1471-8286.2004.00684.x

79. Peakall R, Smouse PE. GENALEX 6: genetic analysis in Excel. Population genetic software for teaching and research. Mol Ecol Notes. 2006;6: 288–295. doi:10.1111/j.1471-8286.2005.01155.x

80. R Core Team. R: A Language and Environment for Statistical Computing. Vienna, Austria: R Foundation for Statistical Computing; 2022. Available: https://www.R-project.org/

81. Pinheiro J, Bates D, DebRoy S, Sarkar D, R Core Team. nlme: linear and nonlinear mixed effects models. 2025. Available: https://CRAN.R-project.org/package=nlme

82. Weir BS, Cockerham CC. Estimating F-statistics for the analysis of population structure. Evolution. 1984;38: 1358–1370.

83. Goudet J. FSTAT, a program to estimate and test gene diversities and fixation indices (version 2.9.3). Université de Lausanne, Lausanne, Switzerland; 2001.

84. Goudet J, De Meeüs T, Day AJ, Gliddon CJ. The different levels of population structuring of the dogwhelk, *Nucella lapillus*, along the south Devon coast. In: Beaumont AR, editor. Genetics and Evolution of Aquatic Organisms. London, UK: Chapman & Hall; 1994. pp. 81–95.

85. Krueger-Hadfield SA, Guillemin M-L, Destombe C, Valero M, Stoeckel S. Exploring the genetic consequences of clonality in haplodiplontic taxa. J Hered. 2021;112: 92–107. doi:10.1093/jhered/esaa063

86. Stoeckel S, Arnaud-Haond S, Krueger-Hadfield SA. The combined effect of haplodiplonty and partial clonality on genotypic and genetic diversity in a finite mutating population. J Hered. 2021;112: 78–91. doi:10.1093/jhered/esaa062

87. Brown AHD, Feldman MW, Nevo E. Multilocus structure of natural populations of *Hordeum spontaneum*. Genetics. 1980;96: 523–536. doi:10.1093/genetics/96.2.523

88. Agapow P, Burt A. Indices of multilocus linkage disequilibrium. Mol Ecol Notes. 2001;1: 101–102. doi:10.1046/j.1471-8278.2000.00014.x

89. Kamvar ZN, Tabima JF, Grünwald NJ. *Poppr*LJ: an R package for genetic analysis of populations with clonal, partially clonal, and/or sexual reproduction. PeerJ. 2014;2: e281. doi:10.7717/peerj.281

90. ArnaudLJHaond S, Duarte CM, Alberto F, Serrão EA. Standardizing methods to address clonality in population studies. Mol Ecol. 2007;16: 5115–5139. doi:10.1111/j.1365-294X.2007.03535.x

91. Pritchard JK, Stephens M, Donnelly P. Inference of population structure using multilocus genotype data. Genetics. 2000;155: 945–959. doi:10.1093/genetics/155.2.945

92. Earl DA, vonHoldt BM. STRUCTURE HARVESTER: a website and program for visualizing STRUCTURE output and implementing the Evanno method. Conservation Genet Resour. 2012;4: 359–361. doi:10.1007/s12686-011-9548-7

93. Jakobsson M, Rosenberg NA. CLUMPP: a cluster matching and permutation program for dealing with label switching and multimodality in analysis of population structure. Bioinformatics. 2007;23: 1801–1806. doi:10.1093/bioinformatics/btm233

94. Rosenberg NA. DISTRUCTLJ: a program for the graphical display of population structure. Mol Ecol Notes. 2004;4: 137–138. doi:10.1046/j.1471-8286.2003.00566.x

95. Kopelman NM, Mayzel J, Jakobsson M, Rosenberg NA, Mayrose I. CLUMPAK: a program for identifying clustering modes and packaging population structure inferences across *K*. Mol Ecol Resour. 2015;15: 1179–1191. doi:10.1111/1755-0998.12387

96. Lenth RV, Piaskowski J. emmeans: estimated marginal means, aka least-squares means. 2025. Available: https://rvlenth.github.io/emmeans/

97. Underwood AJ. Experiments in ecology: their logical design and interpretation using analysis of variance. Cambridge University Press; 1997.

