## Supplemental Figures for "Life-cycle plasticity enables conditional asexual reproduction in a kelp"

### Supplemental information

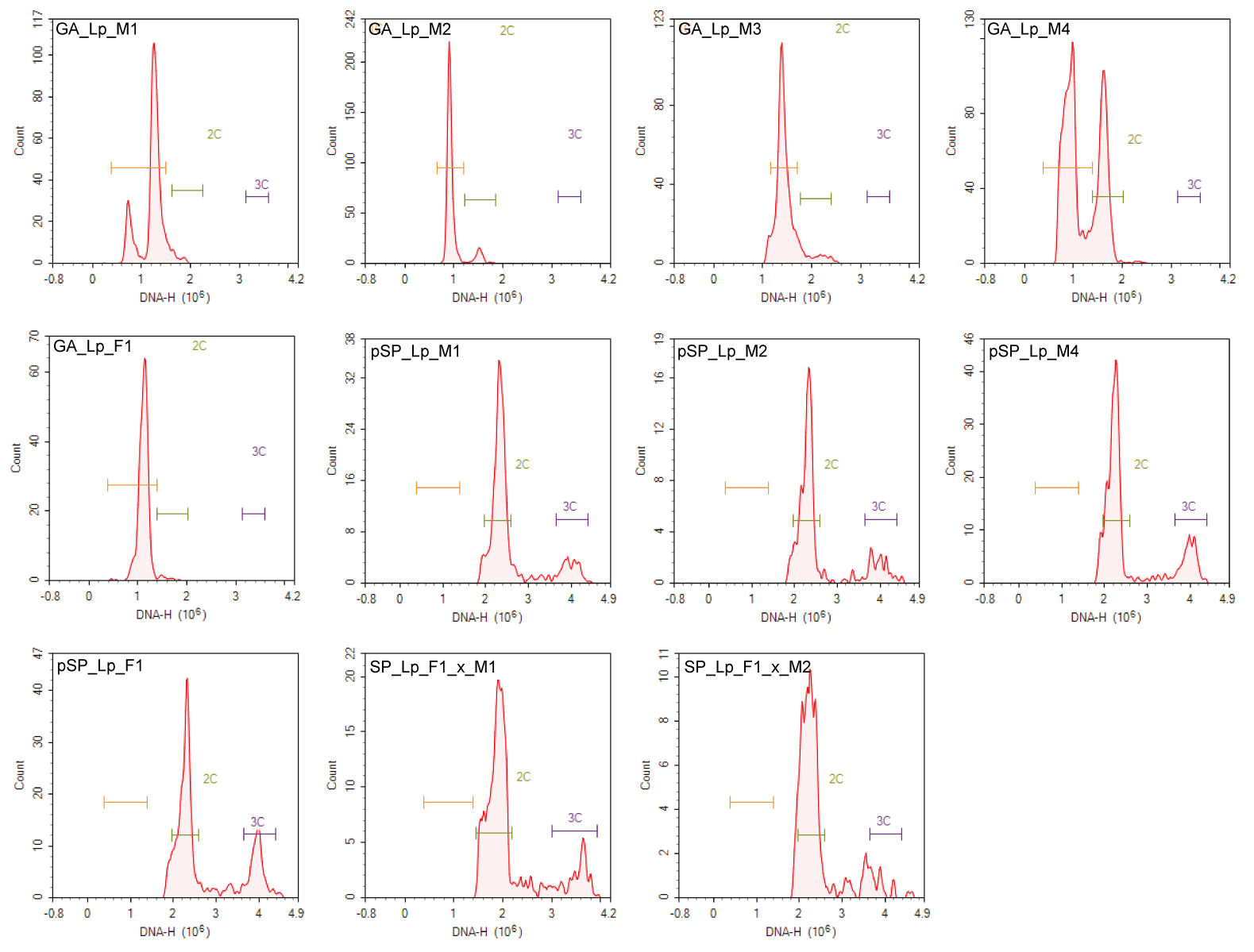

**Figure S1.** Results of flow cytometry analysis to estimate DNA content of *L. pallida* gametophyte (GA), partheno-sporophyte (pSP) and sporophyte (SP) material of five strains (males M1, M2, M3, M4; female F1; cross F1_x_M1, F1_x_M2). Plots show density of fluorescence signals (i.e., number of nuclei) as a function of DNA content (DNA-H).

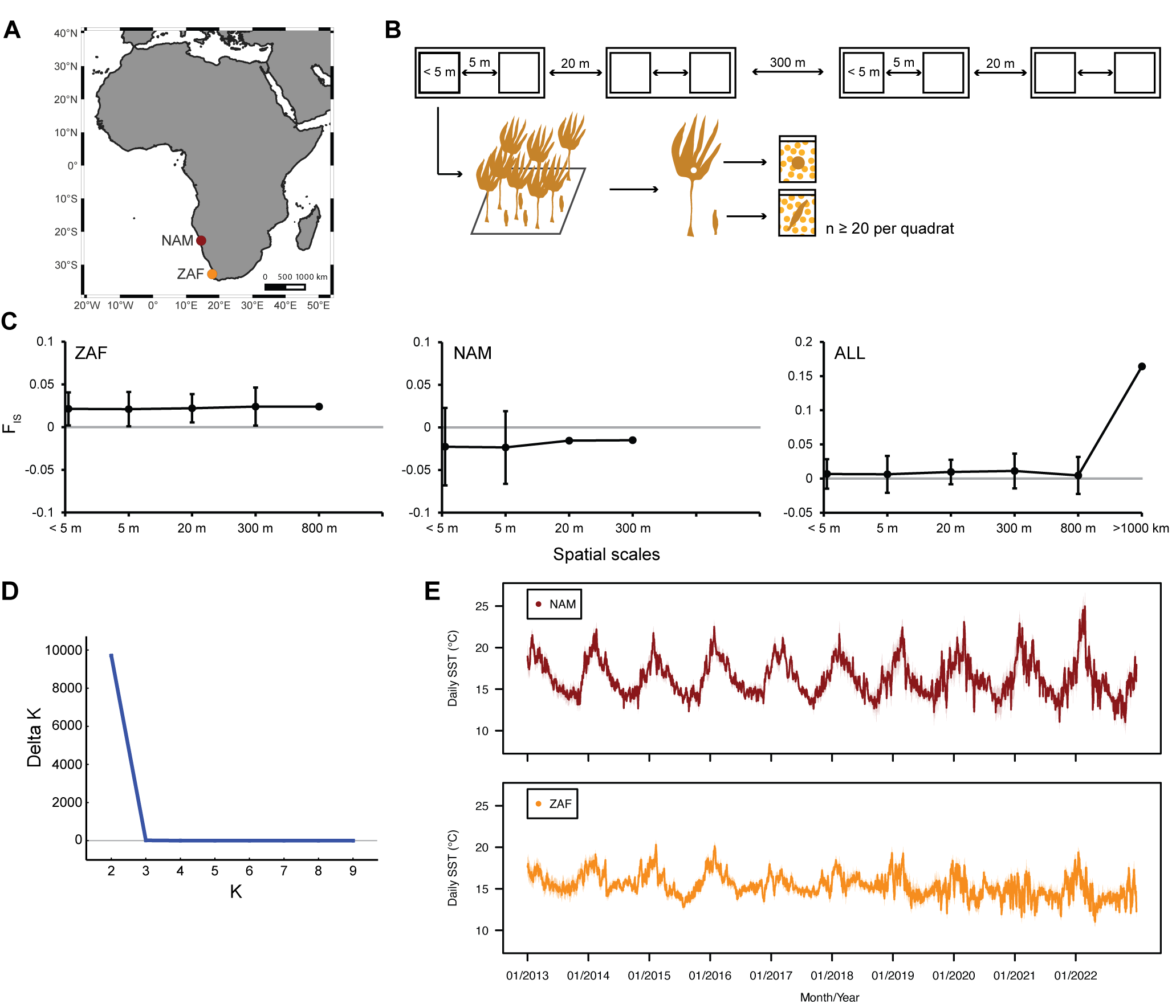

**Figure S2.** Wild populations of *Laminaria pallida* investigated in this study. **(A)** Map depicting the location of Swakopmund, Namibia (NAM) and Paternoster, South Africa (ZAF). **(B)** Schematic hierarchical sampling design applied at the two populations. **(C)** Inbreeding coefficient over spatial scales in the hierarchical sampling design for ZAF, NAM, and both populations (mean ± 95% CI). **(D)** ΔK (Evanno et al., 2005) plotted against number of genetic clusters (K), for K = 2 to K = 9 obtained with Structure Harvester. **(E)** Variation of daily sea surface temperatures for NAM and ZAF from 2013 to 2022 (10 years before time of sampling) obtained from the E.U. Copernicus Marine Service (<https://doi.org/10.48670/moi-00165>) with daily means shown as a solid line and shading indicating standard error.

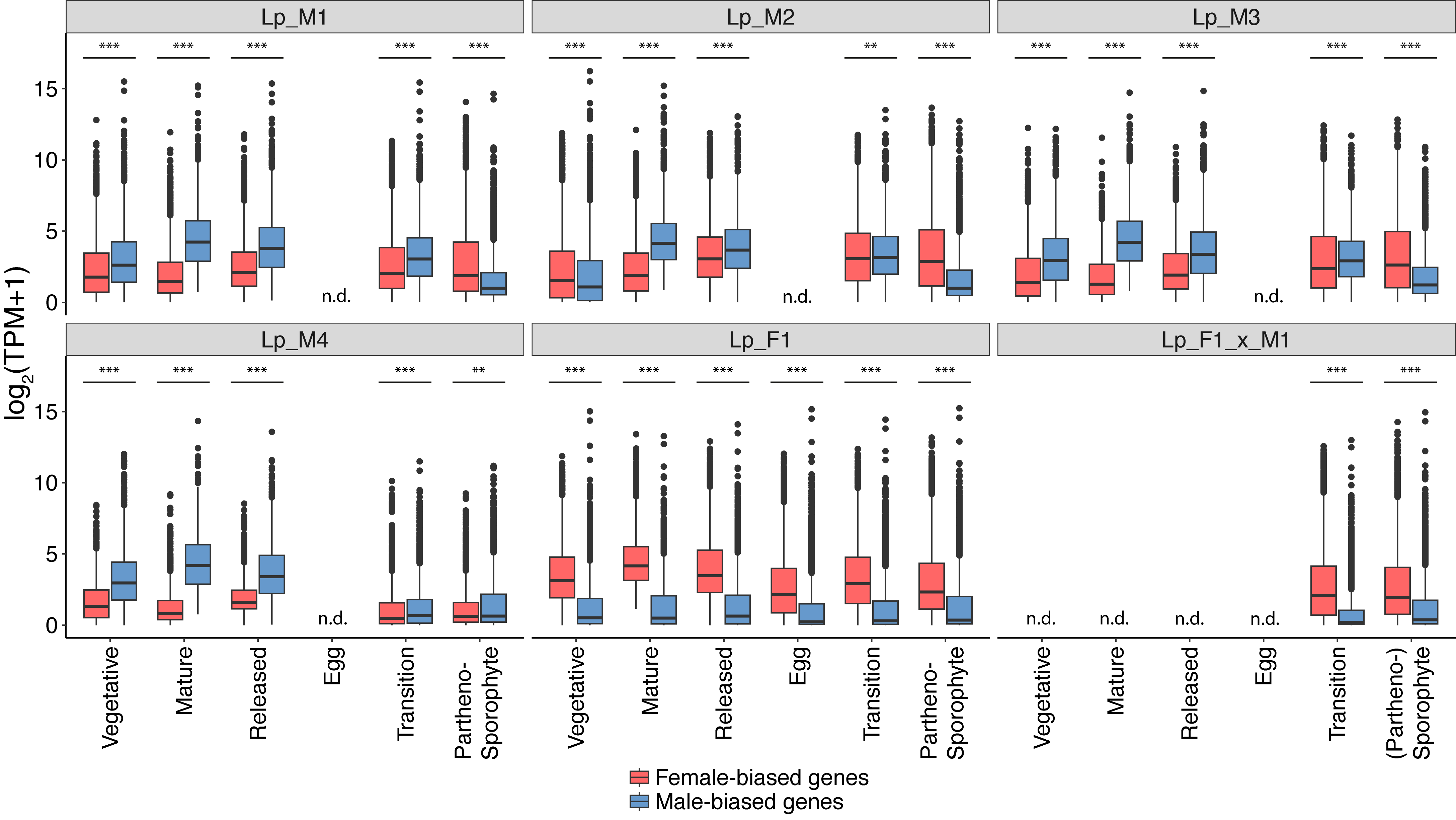

**Figure S3.** Expression of sex-biased genes along development for four male (M1, M2, M3, M4) and one female strain (F1) of *Laminaria pallida* and their cross (F1_x_M1). Significant differences between male- and female-biased gene expression are indicated with asterisks (Wilcoxon rank sum tests; **, p<0.01; ***, p<0.001); n.d., no data.

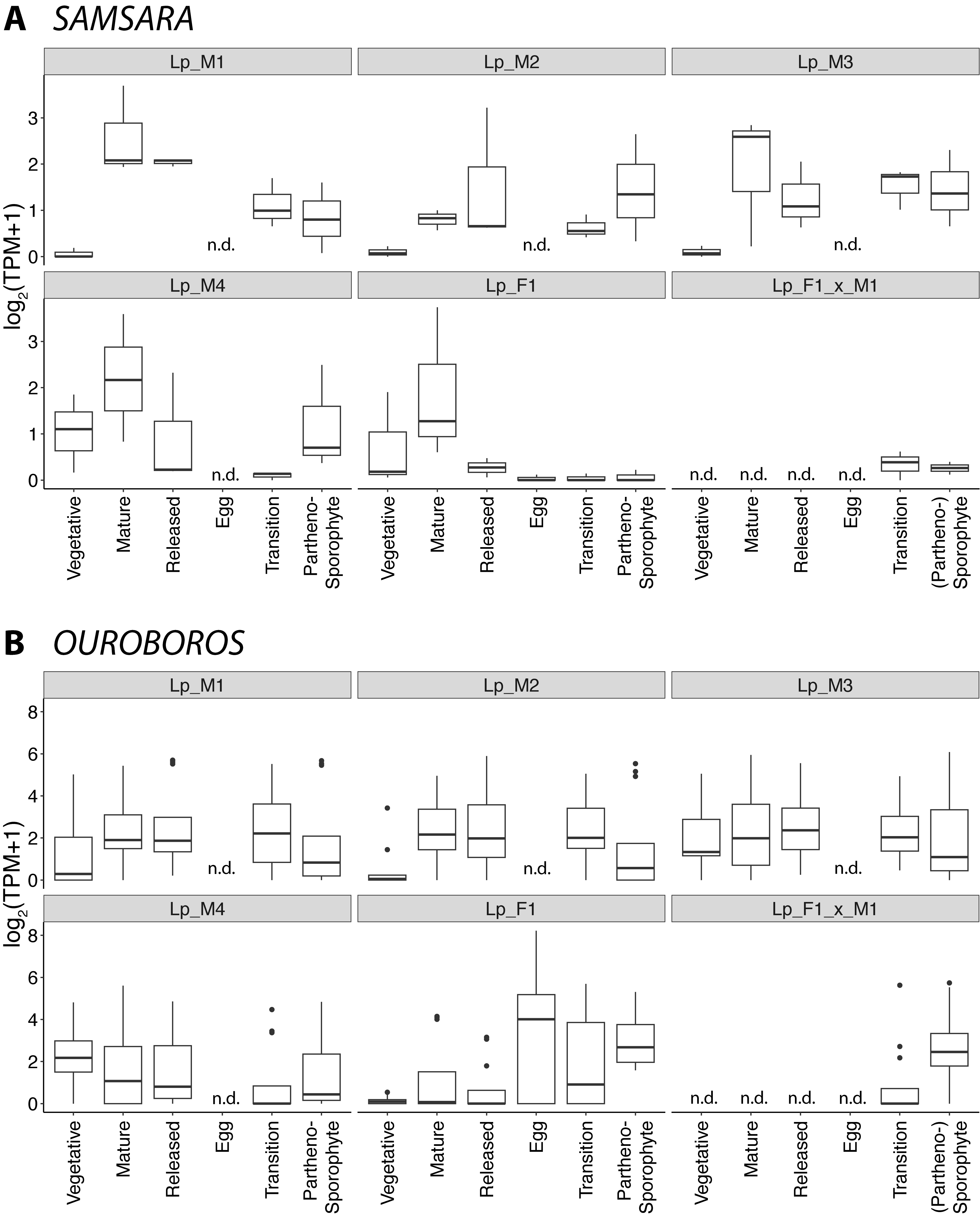

**Figure S4.** Expression of life-cycle related TALE homeodomain transcription factors (A) SAMSARA (1 transcript in n=3 replicates) and (B) OUROBOROS (4 transcripts in n=3 replicates) along development for four male (M1, M2, M3, M4) and one female strain (F1) of *Laminaria pallida* and their cross (F1_x_M1); n.d., no data.

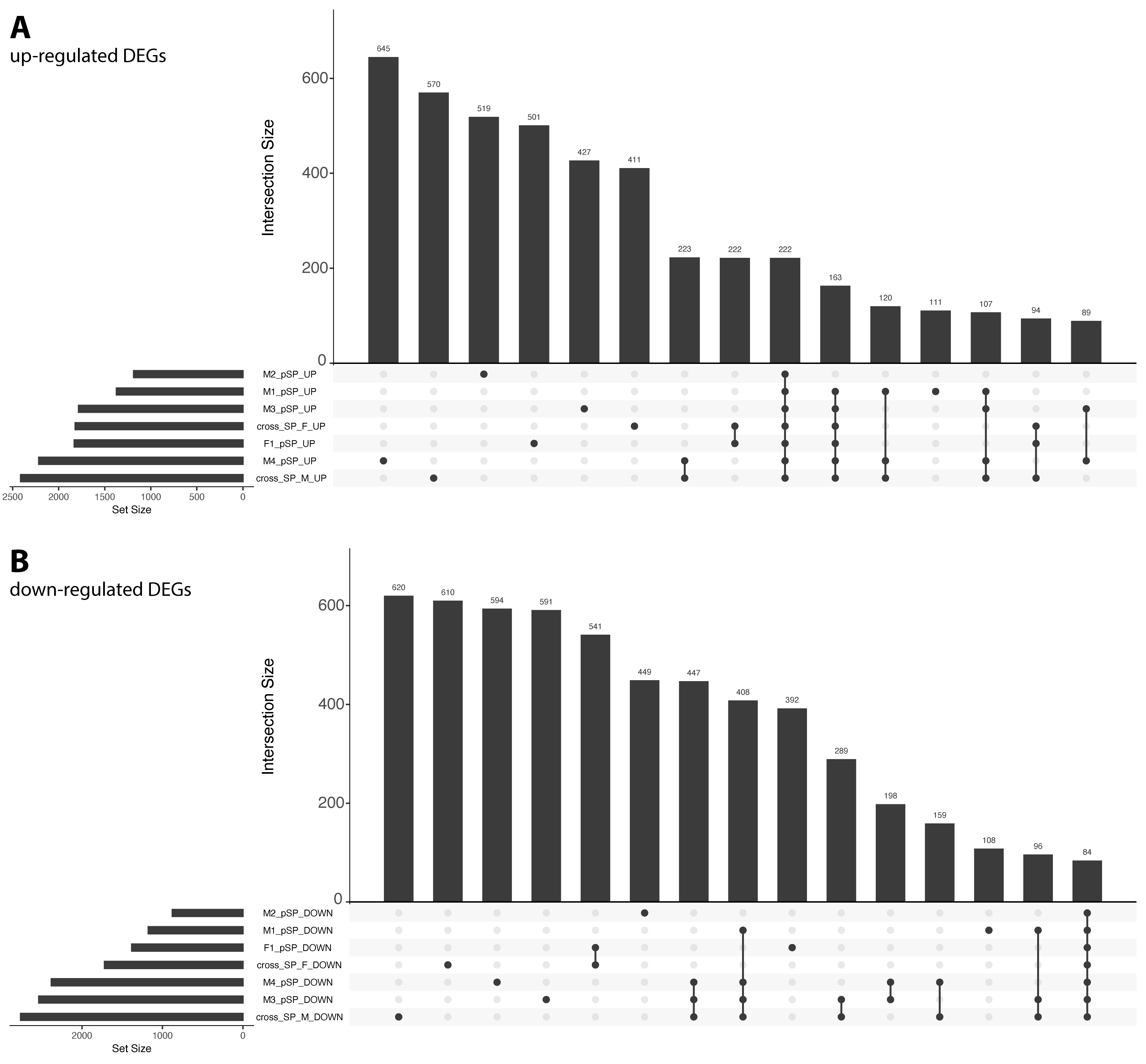

**Figure S5.** Intersection of **(A)** up-regulated and **(B)** down-regulated differentially expressed genes (DEGs) between (partheno-)sporophytes and their corresponding gametophyte for four male (M1, M2, M3, M4) and one female strain (F1) of *Laminaria pallida* and their cross (F1_x_M1). Only the 15 largest intersections are shown.

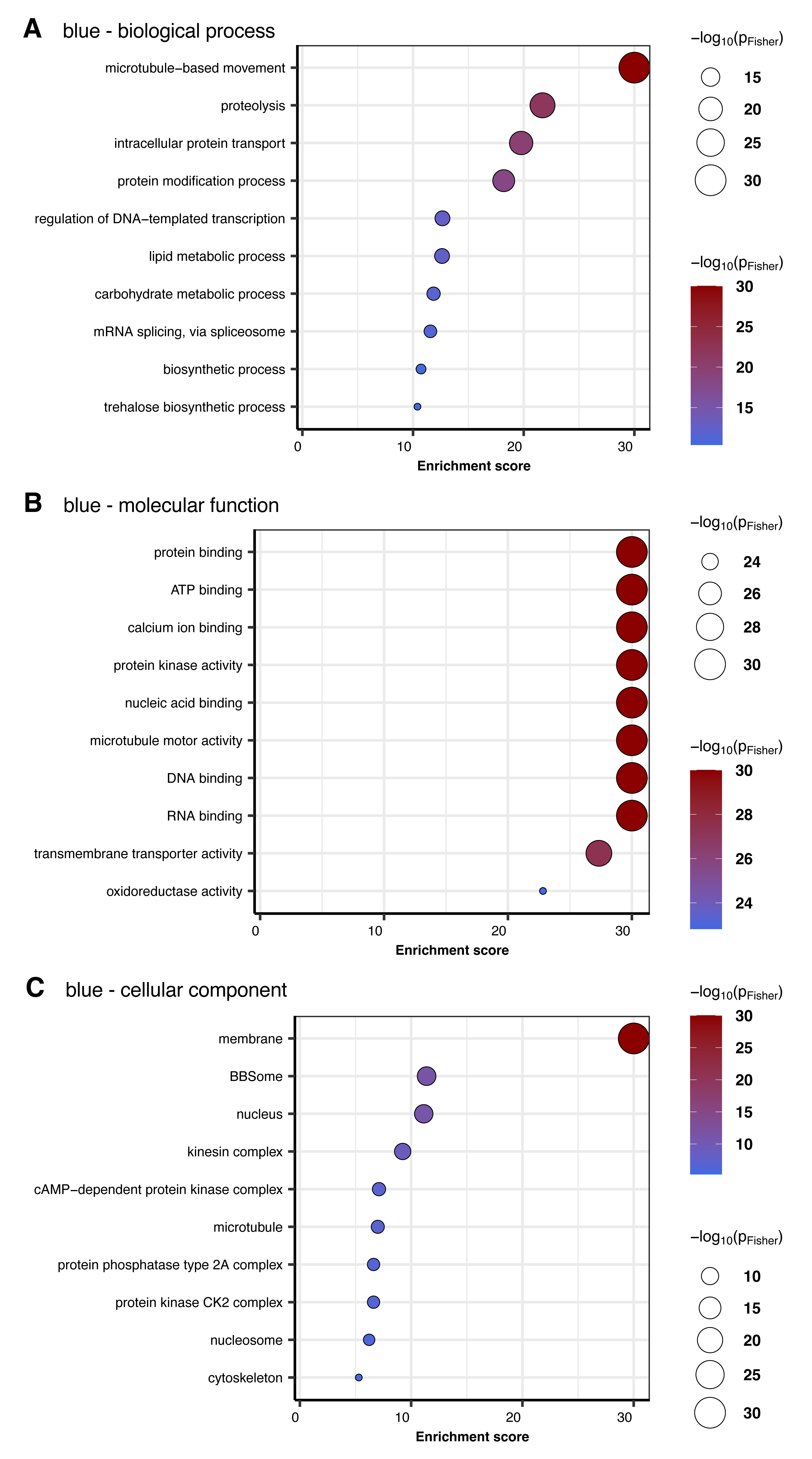

**Figure S6.** Enriched gene ontology (GO) terms for **(A)** biological process, **(B)** molecular function, **(C)** cellular component in module “blue” of weighted gene co-expression network analysis of *Laminaria pallida* development. A maximum of ten most enriched terms is displayed in dotplots sorted by significance (Fisher’s exact test).

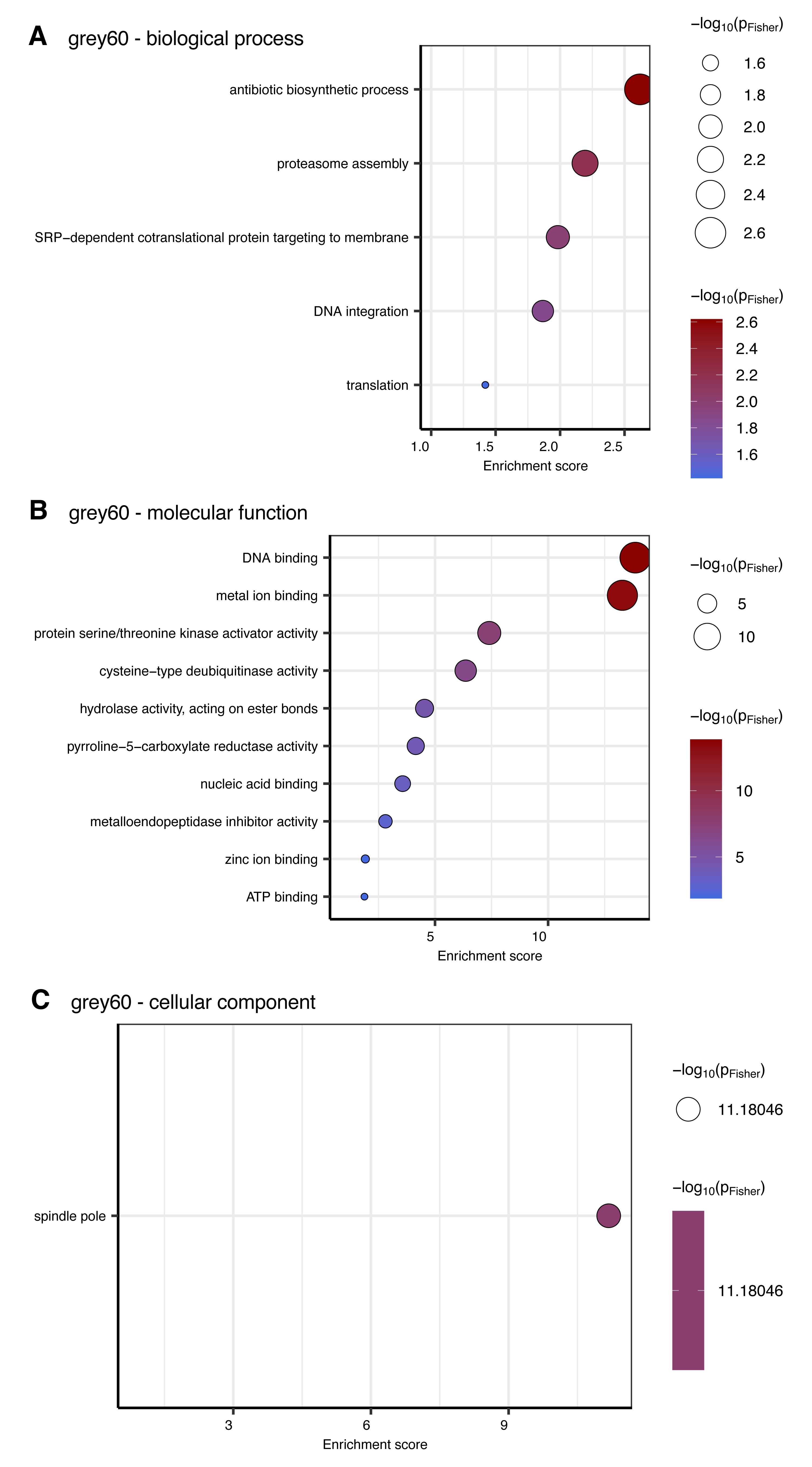

**Figure S7.** Enriched gene ontology (GO) terms for **(A)** biological process, **(B)** molecular function, **(C)** cellular component in module “grey60” of weighted gene co-expression network analysis of *Laminaria pallida* development. A maximum of ten most enriched terms is displayed in dotplots sorted by significance (Fisher’s exact test).

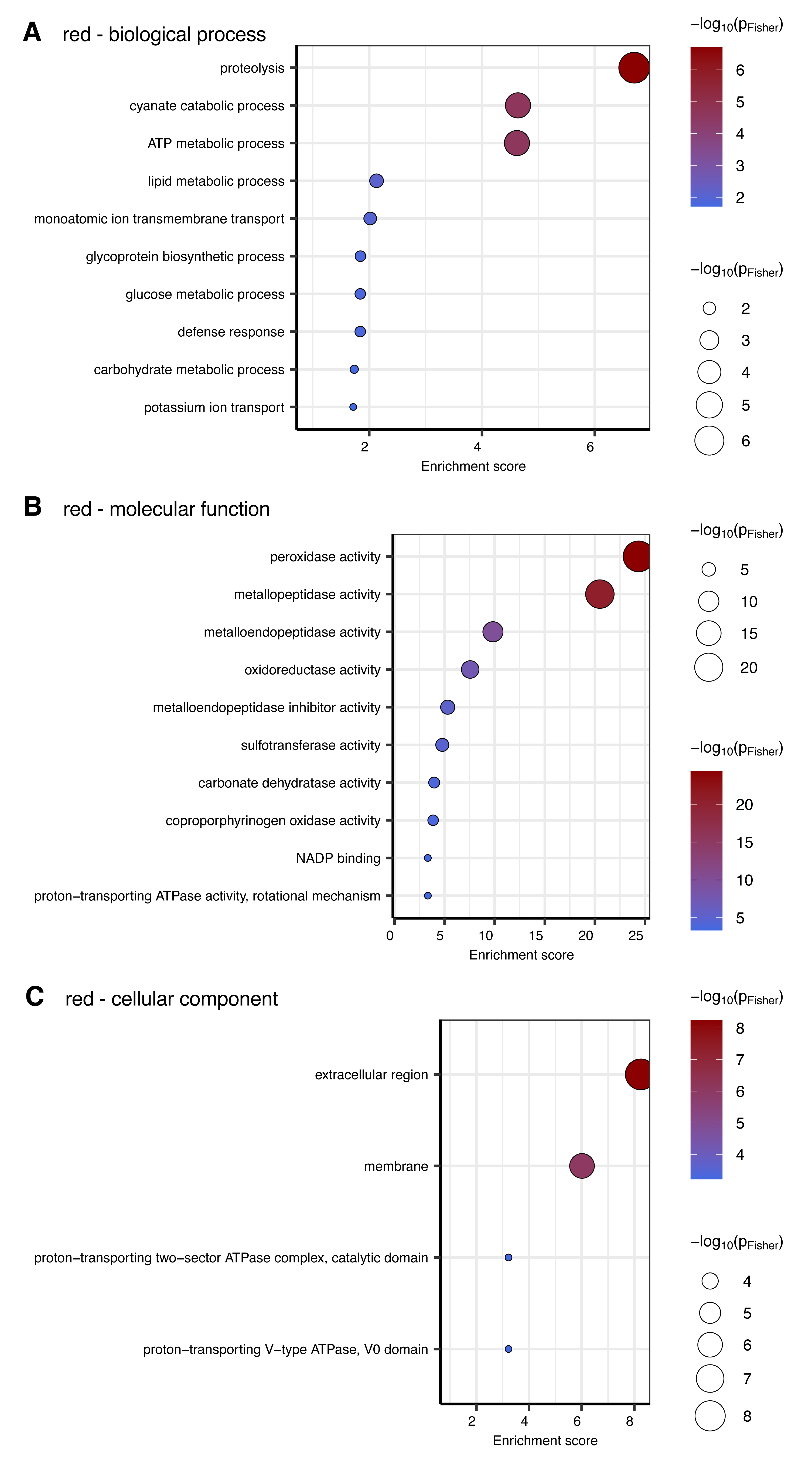

**Figure S8.** Enriched gene ontology (GO) terms for **(A)** biological process, **(B)** molecular function, **(C)** cellular component in module “red” of weighted gene co-expression network analysis of *Laminaria pallida* development. A maximum of ten most enriched terms is displayed in dotplots sorted by significance (Fisher’s exact test).

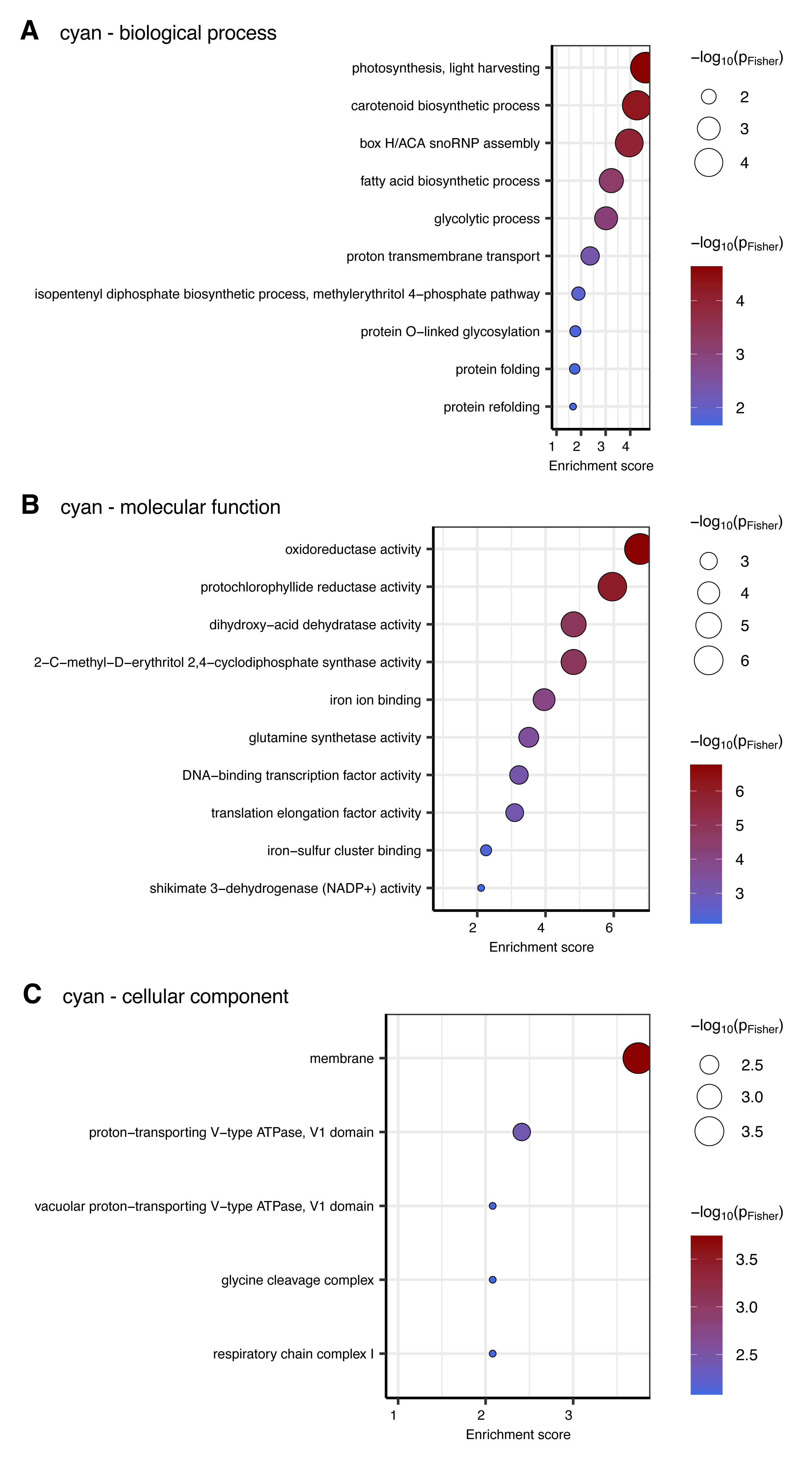

**Figure S9.** Enriched gene ontology (GO) terms for **(A)** biological process, **(B)** molecular function, **(C)** cellular component in module “cyan” of weighted gene co-expression network analysis of *Laminaria pallida* development. A maximum of ten most enriched terms is displayed in dotplots sorted by significance (Fisher’s exact test).
